# Short-wavelength violet light (420nm) stimulates melanopsin-dependent acute alertness responses in zebrafish

**DOI:** 10.1101/825257

**Authors:** Jorge E. Contreras, Thomas S. Lisse, Chifaa Bouzidi, Ann M. Cavanaugh, Anna Matynia, Sandra Rieger

## Abstract

Sunlight throughout the day and seasons strongly influences our biological rhythms and activity. In recent years, it has become evident that night-time overexposure to bright light in urban environments can profoundly affect physiology and behaviour in humans and animals. In particular, the artificial emission of short-wavelength light has been shown to stimulate alertness in humans, but the mechanisms remain largely unknown. Utilising a diurnal larval zebrafish model, we identified instant, non-image-forming (NIF) responses to short-wavelength violet light (~420nm), which are activated only during light exposure, and are reminiscent of alertness, including increased heart rate, enhanced locomotor activity, and pectoral fin beating (for increased oxygen supply). We further determined that these responses are driven by sympathetic neuronal circuits and depend on the zebrafish melanopsin homologue Opn4a. We also found that these responses can be modulated by the sleep-regulatory hormone melatonin, but that melatonin is not essential. Our findings reveal a previously unknown mechanism for violet light-dependent acute alertness.

## Introduction

Diurnal and seasonal differences in available sunlight profoundly influence human and animal physiology and behaviour. Both wakefulness and sleep are sensed by ocular photoreceptors and transmitted via the retinohypothalamic tract to the suprachiasmatic nucleus (SCN) in the hypothalamus ^1–3^. The SCN is the master pacemaker of the circadian clock and synchronizes sleep-wake cycles through communication with local transcription/translation-based feedback oscillator and systemic cues that are present in most tissues ^4^. To maintain proper physiological rhythmicity, the circadian clock must be reset every dawn in a process known as circadian photoentrainment. Prolonged or excessive exposure to light can significantly modulate the internal rhythm. It has become evident in recent years that extensive exposure to broad-spectrum white light, such as emitted from light-emitting diodes (LEDs) and fluorescent light sources in urban environments can increase alertness and decrease sleep duration in animals ^5, 6^. Moreover, prolonged nocturnal light exposure disrupts circadian rhythmicity, which may underlie sleep disorders ^7–9^ and diseases like cancer ^10, 11^. Circadian rhythmicity genes, such as *Clock* have been identified in invertebrates, including plants, fungi and *Drosophila ^12–14^* and vertebrates, and are members of the bHLH–PAS family of transcription. Originally identified in mice ^15^, homologues genes have also been discovered in invertebrates, such as Drosophila ^16^ and in other non-mammalian and mammalian species including zebrafish ^17, 18^, frogs, lizards, hamsters and humans ^15^. Clock and another member of this family BMAL1 are regulated by violet/ blue light-sensitive cryptochromes *Cry1* and *Cry2*, which are found throughout the animal kingdom, including plants ^19, 20^, Drosophila ^21, 22^, zebrafish ^23^, birds ^24^ and mammals ^25, 26^.

Another class of photoreceptors that stimulate non-image-forming (NIF) light responses are intrinsically photosensitive retinal ganglion cells (ipRGCs), a third class of photoreceptor besides rods and cones, expressing the photopigment melanopsin ^27–29^. Originally discovered in *Xenopus laevis ^30^*, melanopsin has been subsequently detected also in many other vertebrates including zebrafish ^31–34^, chicken ^35^, mice ^36–38^, rats ^39^, primates ^40^ and humans ^41^. Studies in humans have provided evidence for the influence of ipRGCs on cognitive functions ^42–45^. For example, a small population of blind, rod and cone-less patients that retained NIF photoreception show increased physiological and cognitive functions when exposed to short-wavelength light ^43^. Experimental evidence in mice shows that ipRGCs send monosynaptic projections to the SCN ^39^ and other areas in the brain, such as the thalamic intergeniculate leaflet (IGL), which indirectly entrains the circadian clock ^46^. SCN neurons project via the paraventricular nucleus to the pineal organ to regulate the production of the sleep-regulatory hormone melatonin. At night, SCN neurons stimulate norepinephrine release and production of melatonin through cAMP ^47^. During the day, light suppresses cAMP production, thereby leading to a reduction in the levels of melatonin ^48^. In particular, blue light has been associated with increased alertness through melanopsin-dependent melatonin suppression. In particular, short-wavelength blue light increases alertness by suppressing plasma melatonin levels and increasing the core body temperature and heart rate ^42, 45, 49^. However, the shorter wavelength violet light (~420 nm) has been shown to more acutely stimulate subjective alertness in humans ^50^. These acute alerting effects cannot be explained by melatonin suppression, which is significantly lower when compared with blue light ^51^. Whether violet light-induced alertness depends on melanopsin is unclear but evidence in mice and zebrafish suggests that melanopsin, although maximally sensitive to blue light with respect to circadian entrainment, can also absorb light in the violet spectrum ^33, 52, 53^. To further investigate the role of violet light in melanopsin activation and alertness, we have established a larval zebrafish model, which is attractive for studying alertness due to its diurnal sleep-wake behaviour, the rapid establishment of a circadian rhythm within 2 days post fertilization (dpf) ^54^, and the ability to form NIF responses ^32, 55^. Similar to *Xenopus* where melanopsin can regulate photic responses in extra-ocular tissues, such as melanophores, iridial myocytes, and deep brain neurons ^30, 41^, also zebrafish melanopsin is found in extra-ocular regions, including the brain and fibroblasts where it was shown to stimulate NIF responses. For example, larval zebrafish exhibit light-seeking behavior in the absence of the eyes in dim light through activation of the melanopsin homologue Opn4a in pre-optic brain regions ^56^. Recently, cultured zebrafish fibroblasts (ZEM-2S) cells were shown to express various melanopsin genes found in this species^34^, leading to cellular synchronization through activation of components of the phosphoinositol pathway ^57^, a pathway that has also been described in invertebrates and other vertebrate cells for phototransduction ^58–61^. Unlike mammals, which have a single melanopsin gene ^37, 41, 55^, the zebrafish genome harbors five melanopsin genes, *opn4a*, *opn4b, opn4.1, opn4xa* and *opn4xb*, which differ in their expression patterns and origins. While in mammals, melanopsin (Opn4) is found in a small subset of retinal ganglion cells ^27, 62^, in the outer segments of cones in the peripheral retina^28^, and has also been described in the retinal pigment epithelium ^29^, zebrafish melanopsin genes are additionally expressed in extra-ocular regions of the brain, such as the anterior and posterior preoptic area, the posterior tuberculum, and the ventral hypothalamus ^32, 33, 56^. Besides the role of Opn4a in light-seeking behaviour, we show that this photopigment also plays an important role in mediating acute alerting responses in larval zebrafish. These acute responses are not induced blue wavelength light but by the shorter wavelength violet light, leading to increased heart rate, oxygen consumption via pectoral fin beating ^63^, and the enhancement of locomotor activity. They are mediated by adrenergic pathways and can be modulated by melatonin but this hormone is not essential. Our study for the first time demonstrates a novel role for violet light in melanopsin activation and the induction of acute alerting responses in zebrafish.

## Results

### Violet light acutely increases the heart rate in larval zebrafish

We serendipitously discovered that exposure of larval zebrafish to short-wavelength light noticeably increased the heart rate. To identify the specific wavelengths underlying this phenomenon, we projected halogen light through a series of fluorescent filters spanning the visible spectrum (Fig. 1a). This revealed that light in the violet spectrum (410-426 nm) most significantly increased the heart rate (baseline heart rate: 118.7±3.087 bpm vs. violet light: 147.52±4.69 bpm, increase: 28.82±5.62 bpm; standard error of the mean, s.e.m.)(Fig. 1b; Supplementary Movie 1). We also observed an increase in UV (350-400 nm) (increase: 6.81±2.10 bpm; s.e.m), blue light (450-490 nm) (increase: 13.55±1.58 bpm; s.e.m), and green light (515-580 nm) (increase 6.33±1.16 bpm; s.e.m) light, but this increase was significantly lower as compared with violet light. We next generated Intensity response curves (IRCs) for the heart rate response using the UV, violet, blue and green filter set (Fig. 1c). We found that the heart rate in violet light at the lowest comparable energy output level (~3.5 mW/cm^2^) was 2.3-fold higher than in UV light and 4.6-fold higher than in blue light. Green light at ~20 mW/cm^2^ stimulated a heart rate increase that was 5.5-fold lower than in violet light at a similar energy output. Even at the highest energy levels measured in green light (72.4 mW/cm^2^), the heart rate increase (16.8±2.24 bpm) was 2.5-fold lower than the increase induced by the highest measurable energy levels of violet light (26.1 mW/cm^2^). Interestingly, we found that higher energy output levels in violet light induced irregular beating of the heart in zebrafish larvae, reminiscent of a conduction block leading to skipping of the heartbeat. This precluded us from testing a saturating illumination-response range. Nonetheless, our data show that while the maximal increase in the heart rate response is induced by violet light, there is a smaller induction by blue light and UV.

**Figure 1.**
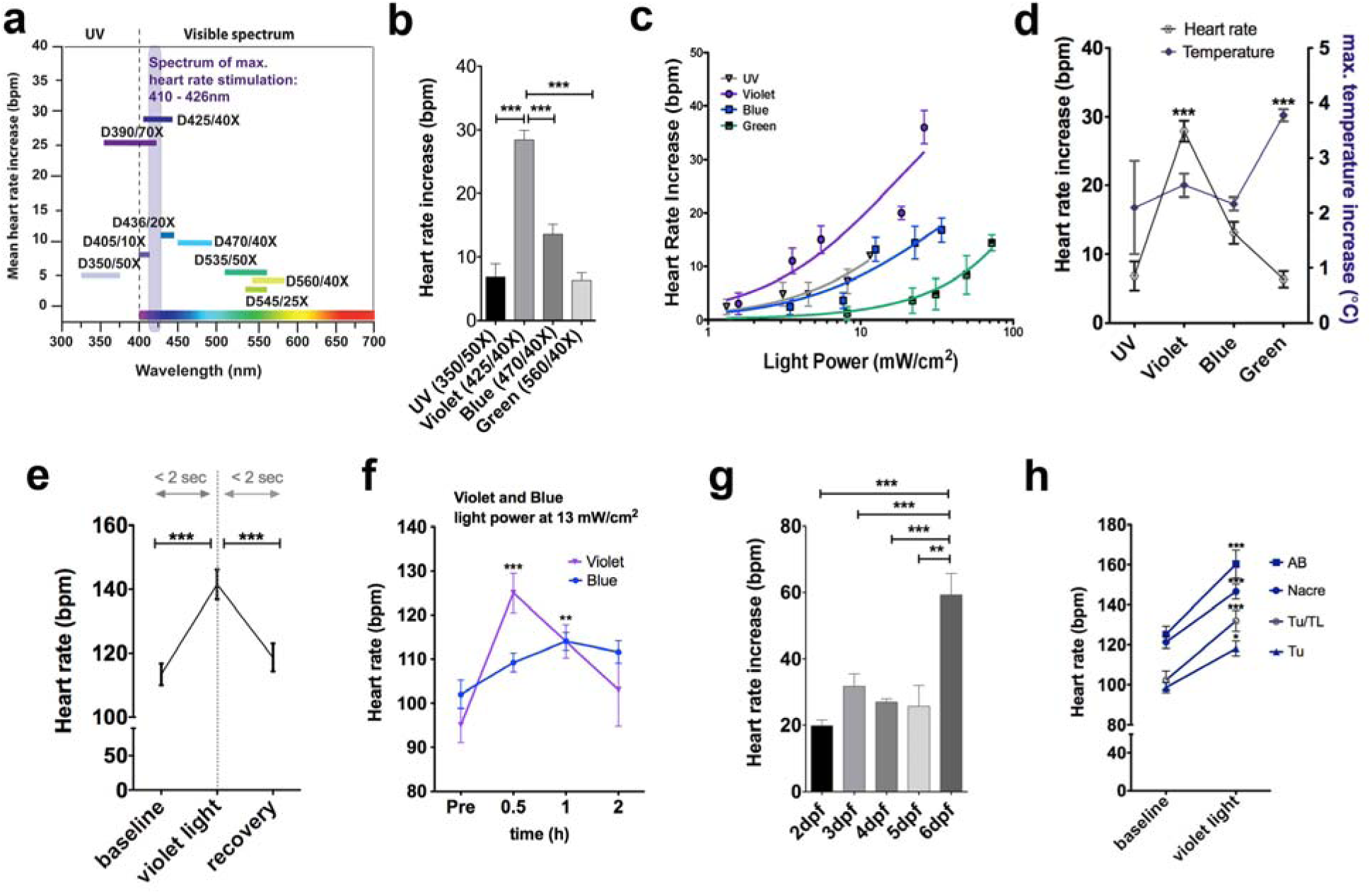
Violet light acutely stimulates a heart rate increase. (**a, b**) Violet light in the range between 410-426 nm maximally increases the heart rate as compared to other wavelengths (12-105 animals per group, s.e.m.). **(c)** Intensity response curves for heart rate response using the UV, violet, blue and green filter set (n=5-6 animals per group, s.e.m). **(d)** The heart rate in UV, violet, blue, and green light is shown in comparison to the wavelength-specific temperature differences (9-157 animals per group, s.e.m). **(e)** The heart rate rapidly increases (within ≤ 2 s) and recovers to baseline levels within a similar time period after violet light exposure (35 animals per group, s.e.m). **(f)** Temporal comparison of the heart rate increase in blue and violet light shows a significant increase from the basal heart rate in each group for both light conditions but with different kinetics and rates (n=6-10 animals per group, s.e.m). **(g)** The violet light-induced heart rate increase can be measured as early as 2 dpf but is more significant in older animals. **(h)** Genetic background does not affect the violet light-induced heart increase (n=12 animals per group, s.e.m). Asterisks above groups represent comparisons to the first group. Brackets indicate alternative comparisons. Statistical comparisons were made using One-Way ANOVA. *P*<0.05, P**<0.01, ***P<0.001*

To rule out that a rise in temperature in the violet spectrum stimulates the heart rate, we compared the temperatures at the plane of the specimen in violet, UV, blue and green light. Since only violet light significantly stimulates the heart rate, we hypothesised that the temperature should be highest in this spectrum. Consistent with the high energy output levels observed in green light, we found that the temperature measured over the time course of 15 sec (which is the maximum time in which we analysed the heart rate) was most significant in green light (mean: +3.77±0.10 °C) as compared with violet light (mean: +2.5±0.21 °C), UV (mean: 2.1±0.84 °C) and blue light (mean: 2.16±0.12 °C)(Fig. 1d). Thus, the heart rate increase observed in violet light cannot be explained by a rise in temperature.

We next assessed the kinetics of the heart rate increase in violet light by measuring the onset and offset of the response. We found that a maximal heart rate increase was observed within only ≤ 2 s, which was reverted to baseline levels at a similar rate upon termination of violet light exposure (Fig. 1e). Blue light has been shown to stimulate alertness in humans ^42, 43, 45^ but subjective alertness ratings showed a more acute effect for violet light ^50^. Given that we did not observe a significant heart rate increase in blue light, we wondered whether blue light requires longer exposure to increase the heart rate. To test this, we compared the heart rate increase in blue light to that of violet light over the time course of 2 hours. We observed that violet light at 13 mW/cm^2^ had a strong heart rate stimulatory effect during the first 30 minutes of continuous exposure (pre: 95±3.92 vs. 30 min: 125±4.49) but the heart rate declined close to pre-exposure levels (103±8.25) after continuous exposure for up to 2 hours (Fig. 1f). In contrast, zebrafish exposed to blue light under the same conditions moderately but steadily increased the heart rate, which became most significant after 1 hour (pre: 102±3.22; 30 min: 109.2±2.15; 1 h: 114±2.0; 2 h: 111.6±2.56) and at this time point reached a similar level as the heart rate of violet light exposed animals. This suggests that violet light induces the heart rate more acutely and transiently than blue light.

Because we observed that the heart rate increase was attenuated when we anaesthetised the larvae with 0.4 mM of the sodium channel blocker tricaine (MS-222) (control: 22.00±5.40 bpm vs. tricaine: 10.11±4.22 bpm; standard deviation, s.d.)(Supplementary Fig. 1), we generally performed all experiments with unanesthetized larvae mounted in 1.2 % low-melt agarose. The violet light response was first detectable at 2 dpf, which coincides with the initiation of visual and cardiovascular system maturation in zebrafish ^64, 65^, and the establishment of a circadian melatonin rhythm ^54^. Consistent with this, we found that this response was potentiated in older larvae at a point when both systems are more matured (2 dpf: 19.79±1.80 bpm vs. 6 dpf: 59.26±6.52 bpm; s.e.m)(Fig. 1g). We next determined the heart rate response in different strains to account for genetic background variations. We analysed AB, Tuebingen (Tu), Tu/Tupfel/Longfin (Tu/TL), and Nacre strains and found that all strains showed a significant heart rate increase in violet light (AB: + 30.95±1.30 bpm, Tu: + 19.83±1.83 bpm, Tu/TL: + 32.72±2.50 bpm, Nacre: + 30.95±1.30 bpm; s.e.m)(Fig. 1h), suggesting that this response is independent of genetic background. Taken together, these findings suggest that violet light stimulates NIF responses in larval zebrafish.

### The heart rate increase depends on ocular photo-transduction

We next sought to investigate whether the heart rate increase is mediated by ocular or extra-ocular photoreception. While NIF responses in mammals are mediated by ipRGCs, in zebrafish extra-ocular tissues can also mediate NIF responses. To assess this, we surgically transected the optic nerves adjacent to the eyes in 3 dpf larvae using glass capillary needles (Supplementary Figs. 2a, b, b’). This manipulation significantly attenuated the heart rate response to violet light (increase prior to transection: 48.63±3.35 bpm vs. after transection: 15.82±2.88 bpm; s.e.m; n=5)(Supplementary Fig. 2c). Because traction applied to extraocular muscles and compression of the eyeball due to our surgery may have induced an oculocardiac reflex, which is mediated by activation of the ophthalmic branch of the trigeminal cranial nerve and the vagus nerve of the parasympathetic nervous system, leading to a lower heart rate, we also utilized a genetic ablation system. We utilized the nitroreductase cell ablation system ^66, 67^ to generate transgenic animals expressing the bacterial enzyme nitroreductase (NTR) encoded by the *nfsB* gene under the *isl2b* promoter (Tg(*isl2b:nfsB-mCherry*))(Fig. 2a and Supplementary Fig. 3a, a’). This promoter was shown to activate expression in RGCs and somatosensory neurons of larval zebrafish ^68^. RGC and somatosensory neurons were ablated at 3 dpf with metronidazole (MTZ), which is converted by NTR into a cytotoxin, leading to cellular death within 24 hours. When starting treatment at 2 dpf, we observed a significant loss of most RGCs and their projections by 3 dpf (Fig. 2a’). Analysis of the heart rate in these animals revealed a significant reduction in the response to violet light (3 dpf: DMSO control: 32.40±2.40 bpm vs. ablated: 9.33±1.45 bpm; 4 dpf: DMSO control: 26.73±1.30 bpm vs. ablated: 14.36±1.57 bpm; s.e.m) (Fig. 2b). In contrast, MTZ treatment of non-NTR expressing transgenic Tg(*isl2b*:GFP) ^68^ control larvae did not attenuate the heart rate increase induced by violet light (untreated: 20.85±5.77 bpm vs. ablated: 22.87±8.99 bpm; s.d.)(Supplementary Figs. 3b-d).

**Figure 2.**
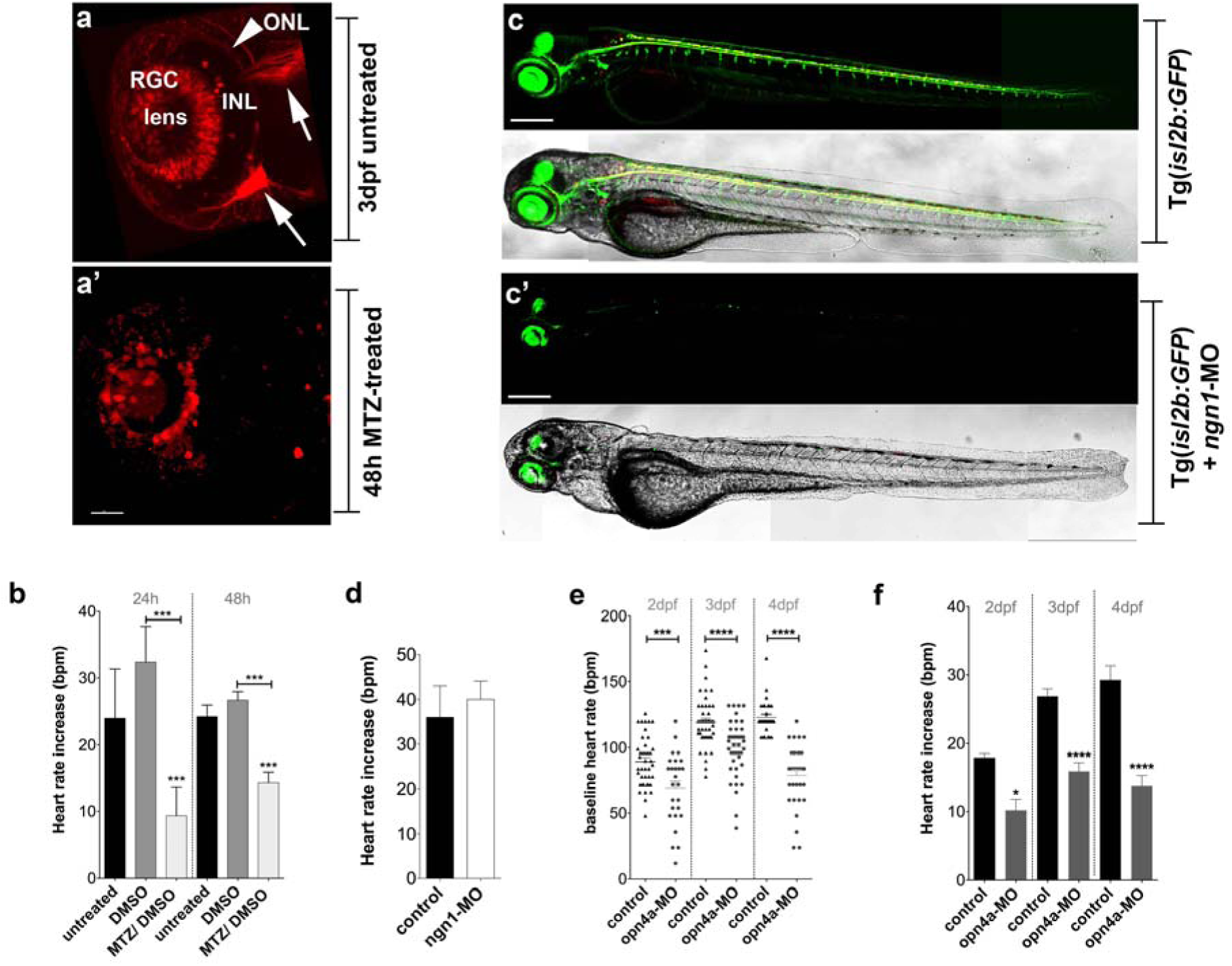
Violet light stimulates the heart rate through melanopsin (*opn4a*) (**a, a’**) Eye of a transgenic Tg(*isl2b:nfsB-mCherry*) control (a) and a MTZ-treated (a’) larva at 4 dpf. (a) The top arrow points to retinal ganglion cell (RGC) fibres that project into the optic tectum, and the bottom arrow points to the trigeminal ganglion. The arrowhead highlights the outer nuclear layer (ONL). (a’) Most neurons and their projections are absent after 48 h of MTZ treatment. Scale bar: 40 µm. (**b**) Comparison of the heart rates in the untreated, DMSO-treated, and RGC-ablated (MTZ-treated) larvae (9-20 animals per group, s.e.m) shows that the violet light-induced heart rate increase is attenuated in the RGC-ablated larvae. (**c, c’**) Fluorescent RGCs and sensory neurons in the Tg(*isl2b:GFP*) wild-type (c) and *ngn1*-knockdown larvae (c’). (c’) GFP^+^ RGCs and their projections are present despite the absence of somatosensory neurons. Scale bars: 250 µm. **(d)** Sensory neuron-deficient *ngn1*-MO larvae show a similar heart rate increase compared with the controls when exposed to violet light (9 animals per group, s.d.). **(e)** Comparison of the baseline heart rates in the control and *opn4a*-atg morphants at 2 dpf, 3 dpf, and 4 dpf (26-48 animals per group, s.e.m) reveals a significantly lower baseline heart rate in the *opn4a*-atg morphants. **(f)** Comparison of violet light-induced heart rate increase in control and *opn4a*-atg morphants at 2 dpf, 3 dpf, and 4 dpf reveals a significantly attenuated response in the *opn4a*-atg morphants (9-25 animals per group, s.e.m). Asterisks above columns represent comparisons to the first column in each group. Brackets indicate alternative comparisons. Statistical comparisons were made using One-Way ANOVA (b, e, f) and Student’s t-test (d). *\*P<0.05, ***P<0.001, ****P<0.0001*

**Figure 3.**
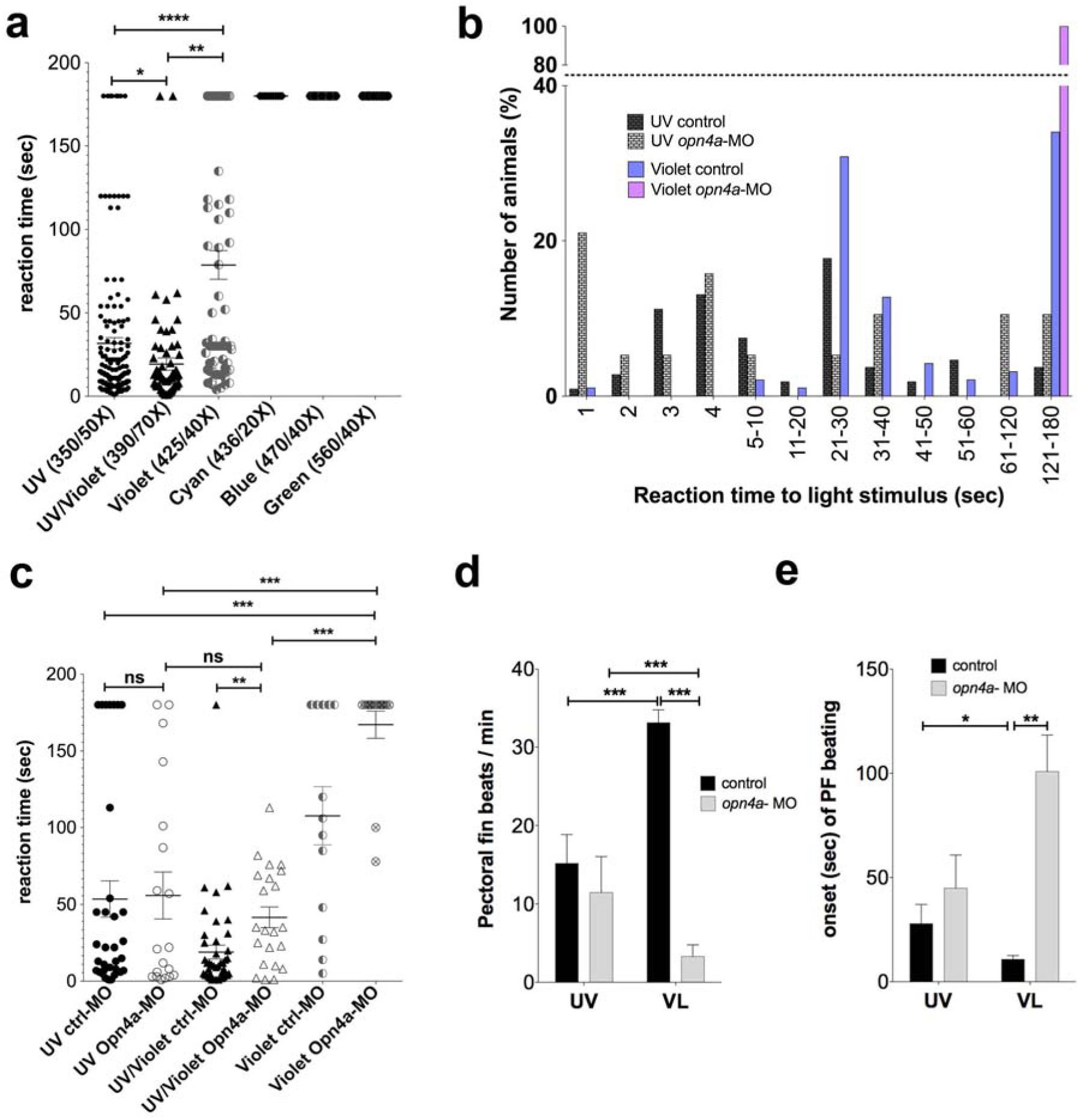
Violet light stimulates alertness-like behavioural responses in zebrafish. **(a)** The onset of the locomotor response at 3 dpf in UV/VL (n=74) is significantly faster than in UV light alone (n=167). Violet light elicits a significantly delayed response (n=72). No response is observed in cyan, blue, and green light at this age (10-16 animals, s.e.m). **(b)** Distributions of locomotor response onsets at 3 dpf for wildtype and *opn4a*-morphants in UV light and violet light. Shown is the percentage of total larvae assessed within each group. The data was normalized for spontaneous swimming behaviour. **(c)** Locomotor responses at 3 dpf are largely abolished in the *opn4a*-atg morphants exposed to violet light (14 animals per group, s.e.m) but not in UV light. UV/VL elicits a response in the *opn4a*-atg morphants that is more similar to the reaction times in UV light alone (n=22-44 animals per group, s.e.m). **(d)** The rate of pectoral fin beats per minute is significantly higher in violet light as compared with UV light, whereas *opn4a*-spl morphants have a significantly lower rate in violet but not UV light (10-40 animals per group, s.e.m). **(e)** The onset of pectoral fin beating is more rapid in violet light as compared to UV light in the controls, and significantly delayed in the *opn4a*-spl morphants exposed to violet light. In contrast, the UV light response appears to be similar to the controls (maximal measured time was 180 s) (10-25 animals per group, s.e.m). Asterisks above brackets indicate comparisons between groups. Statistical comparisons were made using One-Way ANOVA. *P*<0.05, P**<0.01, ***P<0.001, ****P<0.0001*

Because MTZ ablation in this transgenic line eliminated both RGCs and somatosensory neurons, including trigeminal neurons, we wanted to exclude that ablation of somatosensory neurons did not causes the attenuation of the violet light-induced heart rate increase. We therefore generated RGC-intact but somatosensory neuron-deficient zebrafish larvae by knocking down the bHLH transcription factor *neurogenin* 1 (*ngn1*), which is required for somatosensory neuron specification ^69, 70^. We expected that if somatosensory neurons are critical for this response, we should see a similar attenuation as with the MTZ-mediated ablation. We validated the knockdown efficiency of *ngn*-1 in transgenic Tg(*isl2b*:GFP) larvae in which *ngn1*-positive somatosensory neurons (trigeminal, dorsal root ganglia and Rohon beard neurons) were ablated but *ngn1*-negative, intact RGCs retained GFP-fluorescence (Figs. 2c and c’). Both controls and somatosensory neuron-deficient, RGC-intact larvae showed a similar heart rate increase induced by violet light (WT: 36.00±12.00 bpm vs. *ngn1*-MO: 40.00±9.79 bpm; s.d.)(Fig. 2d), indicating that the response is mediated through ocular photoreception. Taken together, these findings suggest that retinal pathways are involved in the heart rate increase induced by violet light but this response does not require somatosensory innervation of the skin.

### Opsin 1 short-wave sensitive 2 (Opn1sw2) is not essential for the violet light response

We next sought to identify the responsible photopigment, and although an image-forming opsin, the cone pigment *opsin 1, short-wavelength-sensitive 2* (*opn1sw2*) represented the only known opsin with a peak absorbance in the violet spectrum at ~416 nm ^71^. Also its upregulation in the zebrafish retina by 50 hours post fertilization ^72^ coincided with our observations. To test for a possible involvement of this photopigment, we generated two *opn1sw2* morpholinos to knockdown its expression. We first tested a translation blocking morpholino by injecting 1-10 ng into fertilized eggs. This MO however caused severe apoptosis in the forebrain of embryos at 1 dpf, from which the embryos never recovered, even when co-injected with a *p53* morpholino (not shown). We therefore also tested a splice MO targeting the exon 1/ intron 1 splice junction. Reverse-transcription PCR showed that the resulting transcript was largely degraded, likely due to nonsense-mediated RNA decay (Supplementary Fig. 4a), which we confirmed with quantitative PCR (Supplementary Fig. 4b). Despite the efficient knockdown, the violet light-induced heart rate increase was not significantly altered (Supplementary Fig. 4c). These findings suggest that alternative photopigments that are sensitive to violet light must mediate the heart rate response.

**Figure 4.**
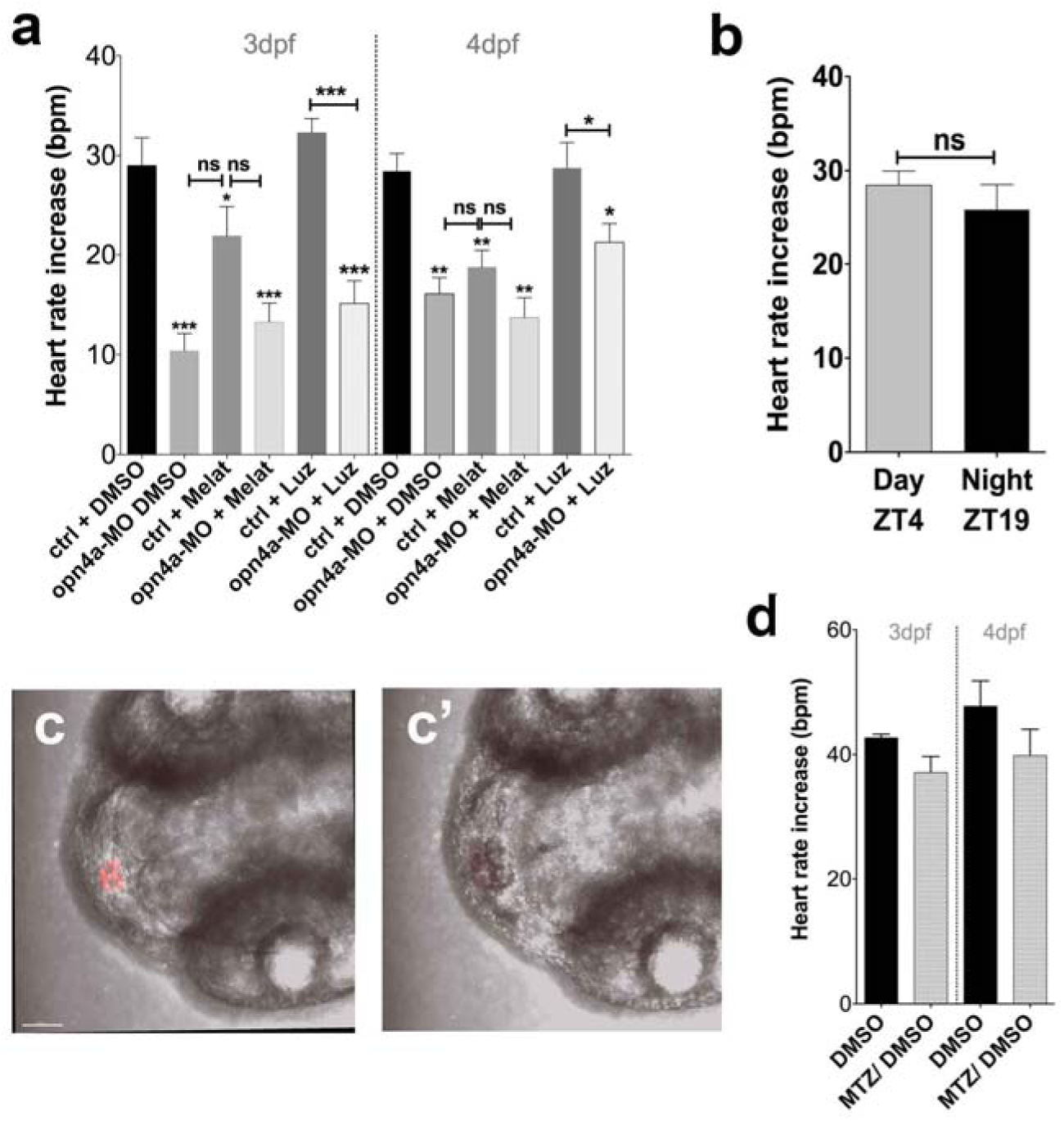
Melatonin can modulate the violet light-induced heart rate increase. **(a)** Comparison of control and *opn4a* (atg and spl) morphant larvae at 3 and 4 dpf treated with DMSO, melatonin (50 µM) and Luzindole (10 µM). Pharmacological treatment with melatonin during daytime (Zeitgeber (ZT) 4-6) attenuates the heart rate increase in the wildtype larvae but does not further attenuate the increase in the *opn4a* morphant larvae. Luzindole-treated control and *opn4a* morphant larvae at 3 dpf do not show any difference to the respective DMSO-treated controls, whereas a slight increase in the heart rate is observed in *opn4a* morphants at 4 dpf (12 - 42 animals per group, s.e.m). **(b)** Comparison of the heart rate increase induced by violet light between Zeitgeber 4 (daytime) and Zeitgeber 19 (night-time) does not reveal a significant difference (13 larvae per group, s.e.m.) (**c, c’**) The pineal gland (PG) is shown before (c, red fluorescence) and after ablation (c’) in a 3-day old larva. Scale bar: 50 µm. (**d**) The violet light–induced heart rate increase does not differ between the DMSO-treated controls and the MTZ-treated, PG-ablated larvae at 3 dpf and 4 dpf (16 animals per group, s.e.m). Asterisks above columns represent comparisons to the first column at each time point. Brackets indicate alternative comparisons. Statistical comparisons were made using One-Way ANOVA (a, d) and Student’s t-test (b). *P*<0.05, P**<0.01, ***P<0.001*

### The violet light-induced heart rate increase depends on melanopsin Opn4a

We next decided to investigate the role of melanopsins since mammalian Opn4 has been shown to transduce NIF responses, and given evidence from a heterologous cell culture expression study suggesting a sensitivity of mouse melanopsin to 424 nm violet light ^52^. The zebrafish melanopsin homologue *opn4a* shares ~70 % of its sequence within the core region with human melanopsin, *OPN4 ^55^*; we therefore decided to focus on this photopigment. Unlike mammalian OPN4, which is expressed in ipRGCs, zebrafish *opn4a* is expressed initially in the presumptive pre-optic area starting around 1 dpf ^56^, and later also becomes upregulated in bipolar, horizontal and amacrine cells of the inner nuclear layer (INL) ^32, 33^. We designed a translation (*opn4a*-atgMO) and splice blocking MO (*opn4a*-splMO) to assess the role of this photopigment in the violet light-induced heart rate increase. The *opn4a*-splMO targets the exon1/ intron 1 boundary (Supplementary Figure 5a-c), which is predicted to prevent splicing of intron 1 and inclusion of a 7.8-kilobase product. Intron 1 inclusion generates a stop codon at position 255 and truncation of the resulting protein at amino acid position 85 (including 30 aa of intron 1). Following injection of either the *opn4a*-atgMO (Supplementary Fig. 6a, a’, b, b’) or the *opn4a*-splMO, some embryos were born with smaller eyes, which we did not find in the control morphants. Compared with the controls, the *opn4a*–atg morphants had a significantly lower baseline heart rate (Fig. 2e)(2dpf: control: 89.34±2.98 bpm vs. *opn4a*-atgMO: 69.23±5.32 bpm; 3 dpf: control: 119.7±2.62 bpm vs. *opn4a*-splMO: 100.3±3.13 bpm; 4dpf: control: 123.00±2.33 bpm vs. *opn4a*-splMO: 78.97±4.33 bpm; s.e.m). We also observed a lower baseline heart rate in the *opn4a*-spl morphants when compared to uninjected larvae and those injected with a control morpholino (3 dpf: uninjected: 113.4±3.42 bpm vs. mismatch-MO control: 110.7±4.66 bpm vs. *opn4a-*splMO: 86±5.30 bpm; s.e.m.)(Supplementary Fig. 6d). These findings implicate *opn4a* in normal cardiac activity.

**Figure 5.**
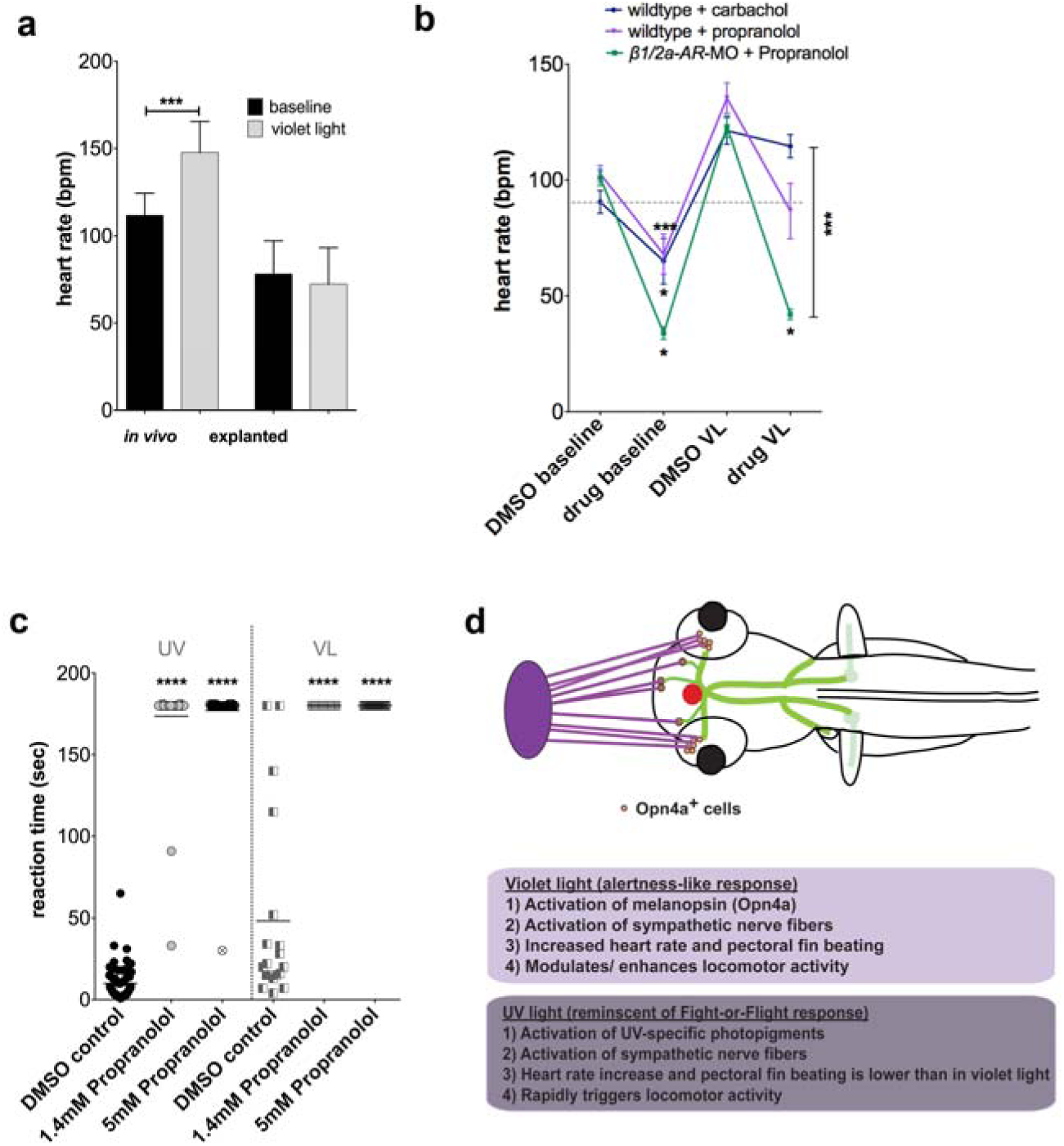
Violet light activates adrenergic receptors to stimulate the heart rate and locomotor activity. **(a)** Explanted hearts (12 animals per group, s.d.) do not respond to violet light in contrast to *in vivo* control hearts (10 animals per group, s.d.). **(b)** Carbachol significantly decreases the baseline heart rate, which can be overcome with violet light (15 animals, s.e.m). Treatment with the β-adrenergic receptor blocker propranolol also lowers the baseline heart rate, but the heart rate is not restored with violet light. This effect is further potentiated in propranolol-treated β*1/2a-AR* double morphants (9 animals, s.e.m). **(c)** UV and violet light-induced locomotor reaction times are significantly decreased in 3 dpf larvae treated with 1.4 and 5 mM propranolol (36 - 45 animals, s.e.m) as compared with control DMSO-treated larvae (19-59 animals, s.e.m). **(d)** Model for light-induced responses: Violet light (indicated by the violet oval and rays) stimulates an alertness-like response mediated through melanopsin (Opn4a) being expressed in ocular and extra-ocular neurons (brown dots) in larval zebrafish. Opn4a activation stimulates sympathetic neuronal circuits (green), leading to an increase in the heart rate and pectoral fin beating (as a means to increase oxygen supply), and the enhancement of basal locomotor activity (upper panel). Melatonin produced in the pineal gland (red) can influence the violet light-induced responses but is not essential. UV light-induced responses are more reminiscent of a fight-or-flight response (lower panel) with rapid locomotor reaction time, and attenuated heart and pectoral fin beating increase. UV light responses similar to violet light responses depend on sympathetic activity. Asterisks in (b, c) indicate comparisons to the control column in each group. Brackets indicate alternative comparisons. Statistical comparisons were made using One-Way ANOVA. *P*<0.05, ***P<0.001, ****P<0.0001*

We next assessed the violet light-induced heart rate increase in *opn4a* morphants and found that the *opn4a*-atg morphants had a significantly attenuated heart rate increase (2 dpf: control: 17.90±0.60 bpm vs. *opn4a*-atgMOs: 10.20±1.59 bpm; 3 dpf: control: 26.93±1.09 bpm vs. *opn4a*-atgMO: 15.93±1.26 bpm; 4 dpf: control: 29.22±2.10 bpm vs. *opn4a*-atgMO: 13.81±1.52 bpm; s.e.m) (Fig. 2f)(Supplementary Movie S2). We observed a similarly attenuated heart rate increase also in the *opn4a*-spl morphants when analysed at 3 dpf (control: 35.14±3.94 bpm vs. *opn4a*-splMO: 17.56±1.46 bpm; s.e.m.)(Supplementary Fig. 6e). We noticed that the heart rate increase was never completely abolished in the *opn4a* morphants, which could be due to variable knockdown efficiencies between animals. Alternatively, other melanopsin homologues or biological systems could be activated and partially compensate for the loss of *opn4a*.

To further validate the specificity of *opn4a* knockdown, we cloned *opn4a* from 4 dpf larvae, and co-injected the mRNA with the splice morpholino into fertilized eggs. Under these conditions, the mRNA although present in all cells of the embryo, is largely degraded around 2 dpf. We therefore analysed the violet light-induced heart rate increase in 2 dpf larvae. We hypothesized that the heart rate would be rescued when we substitute for the loss of *opn4a*, and indeed, we found that the heart rate increase was largely restored to wildtype levels (control: 15.00±2.04 bpm vs. *opn4a-*splMO: 8.50±1.20 vs. *opn4a*-splMO + ZF*opn4* mRNA: 12.0±1.30, s.e.m.) (Supplementary Fig. 6f). We also performed these rescue experiments with human *OPN4* mRNA but this did not rescue the heart rate increase induced by violet light (Supplementary Fig. 6h). Because human OPN4 has been shown to maximally absorb light between 460 and 480 nm, we also stimulated larvae expressing this photopigment with the 470/40X filter. This however also did not increase the heart rate. Thus, the violet light-induced heart rate increase is either a zebrafish-specific mechanism or human OPN4 requires additional co-factors that are not present in the larval zebrafish.

### Violet light stimulates alertness-like behavioural responses

Given that violet light activates melanopsin, and both have been implicated in light-dependent alerting effects in humans ^42, 50, 51^, we wanted to determine whether the violet light-induced heart rate increase could be part of a more global alertness response in zebrafish. The primary brain centres for regulating arousal and alertness are the reticular formation (RF) in the brainstem, which is phylogenetically one of the oldest brain regions, and the closely associated locus ceruleus. As part of the reticular activating system, RF-specific neuronal circuits are modulated by complex interactions between cholinergic and adrenergic pathways that exhibit synergistic as well as competitive actions to regulate sleep-wake behaviour ^73^. In humans, alertness can be assessed through simple reaction-time tasks in psychomotor vigilance tests, neuropsychological assessments (e.g., low-amplitude electroencephalographic and electrooculographic measurements, slow rolling eye movements, and blink rate), or positron emission tomography (PET) and functional magnetic resonance imaging (fMRI) scanning ^74^. Because these assessments are not feasible in larval zebrafish, we sought to establish simple, but measurable, alternative assays that could reveal the behavioural traits reminiscent of alertness. In humans and other diurnal species, light increases locomotor activity, also known as positive masking ^75, 76^. In mice high illumination levels supress locomotion, also known as negative masking, in which melanopsin has been implicated ^77^. To establish whether violet light promotes locomotor activity through melanopsin in zebrafish larvae, we first wanted to characterize the locomotor activity in UV, UV/violet, violet, cyan (426-446 nm), blue and green light. In this assay, we determined the reaction time until a locomotor response was elicited following light exposure, using a maximal observation period of 180 seconds (Supplementary Fig. 7a). We compared the reaction times between 3 and 5 dpf because of the increased activity at 5 dpf. Exposure to UV light induced a faster reaction time as compared with violet light at both days (3 dpf: UV: 29.06±3.82 s vs. violet light: 78.71±8.5 s)(Fig. 3a) (5 dpf: UV: 2.91±0.51 s vs. violet light: 40.35±10.36 s; s.e.m)(Supplementary Fig. 7b). However, when we utilized a band pass filter that allowed UV and violet light (UV/VL) to pass (390/70X), we observed a significant increase in the reaction time as compared to UV light alone (3 dpf: UV/VL: 19.36±3.649 and 5 dpf: UV/VL: 2.00±0.26 s). Therefore, violet light appears to enhance the locomotor response but is not a strong activator on its own. Other wavelengths, such as cyan and green light did not trigger a locomotor response at 3 dpf. A response was noticeable at 5 dpf in cyan (133.7±22.99 s; s.e.m) and green light (81.50±29.06 s; s.e.m), but surprisingly blue light did not elicit a locomotor response either at 3 dpf (180.0±0 s; s.e.m) or 5 dpf (176.8±3.2 s; s.e.m) within 180 sec. These findings suggest that faster reaction times in zebrafish larvae are achieved toward UV light and violet light enhances the response.

Because of the different reaction times in UV versus violet light, we speculated that *opn4a*-mediated responses are unique to violet light and that UV light–mediated locomotor behaviour depends on other photopigments. To test this hypothesis, we assessed the locomotor response in *opn4a* morphants, following exposure to UV and violet light. As expected, the locomotor response was nearly absent when *opn4a* morphants were exposed to violet light (control: 107.7±19.01 s vs. *opn4a*-atgMO: 172.7±7.28 s; s.e.m)(Fig. 3b, c). In contrast, the *opn4a* morphant larvae reacted in UV light similar to the controls (violet control: 53.50±11.80 s vs. violet *opn4a*-atg MO: 55.79±15.30 s). When *opn4a-atg* morphants were exposed to UV/VL, their locomotor reaction time was however more similar to UV light (UV/VL: 41.55±6.79 s; s.e.m.). We found similar results in the *opn4a*-spl morphants (Supplementary Fig. 7c). These findings suggest that violet light enhances locomotor activity through Opn4a, but it is not a strong activator on its own. The significantly delayed onset in violet light could be explained by an enhancement of basal locomotor activity.

In humans, alertness is characterised by rapid and shallow respiration, which enhances alveolar oxygen exchange ^78^. We therefore wanted to assess whether we can utilise respiration as a measure of alertness. One way to measure oxygen supply in larval zebrafish is by assessing the rate of pectoral fin beating, which has been shown to improve oxygen supply. This is achieved through fluid mixing around the upper body by the pectoral fins, which does not contribute to forward propulsion but serves a respiratory function since beating occurs at significantly higher rates when zebrafish larvae are placed in a medium that contains low oxygen levels ^63^. We assessed the rate of pectoral fin beating and the onset compared between UV and violet light within a time period of 180 sec. We found that the rate of pectoral fin beating in violet light was significantly increased (33.15±1.63 PF bpm; s.e.m) compared with UV light (15.2±3.65 PF bpm)(Fig. 3d), and also the onset of beating was more rapid in violet light (10.82±1.79 s; s.e.m)(Fig. 3e; Supplementary Movie S3) when compared with UV light (27.83±9.19 s; s.e.m)(Supplementary Movie S4). In contrast, blue light did not stimulate pectoral fin beating within the same time period (Supplementary Movie S5). While the onset of fin beating was not significantly different between wildtype and *opn4a*-spl morphants (44.86±15.96 s) when exposed to UV light, the morphants had a significantly slower onset of pectoral fin beating when exposed to violet light (100.86±17.43)(Fig. 3d). Also the rate of fin beating was significantly reduced in violet light (3.3±1.47 PF bpm; s.e.m) but not in UV light (11.46±4.60 PF bpm; s.e.m)(Fig. 3e). We should note that pectoral fin beating was absent in 100 % of the *opn4a* atg-morphants and 50 % of the *opn4a-*spl-morphants while the remaining had a decreased response, which we attribute to variations in the MO injections. Together, these findings suggest that violet light activates *opn4a* to increase oxygen supply and the heart rate, which enhances locomotor activity. The violet light responses are distinct from UV light responses in that UV light does not activate *opn4a*. UV light triggers a more rapid locomotor response, whereas the heart rate and pectoral fin beating are activated at a much lower rate.

### Melatonin can modulate the violet light-induced heart rate response

Plasma melatonin suppression inversely correlates with increased alertness in humans, which can be induced to a different degree by exposure to monochromatic light either at 420 nm or 460 nm ^44, 45, 79, 80^. Whereas humans produce melatonin in both the retina and the pineal organ ^81^, the zebrafish pineal gland (PG) is the primary melatonin source and the pacemaker of the circadian clock ^54^. Photic entrainment and the generation of the circadian rhythm does not require the SCN in zebrafish, since a structure similar to the SCN has not been identified ^82^.

Similar to humans however, the role of melatonin in sleep regulation is conserved ^83^. To identify whether differences in melatonin levels modulate the violet light–induced heart rate increase, we modulated melatonin levels in the larvae using pharmacological approaches at 3 dpf larvae, when a sufficient melatonin rhythm is established ^54, 84^. We treated the larvae with various melatonin concentrations to assess the influence on the heart rate (Supplementary Fig. 8). This showed that the heart rate increase induced by violet light was attenuated when larvae were treated with concentrations of 1 µM melatonin but it was not further reduced above this concentration (DMSO: 29.00±2.74 bpm vs. 0.1 µM melatonin: 28.8±4.8 bpm; 1 µM melatonin: 17.0±3.92 bpm; 500 µM melatonin: 14.4±2.4 bpm; s.e.m). A concentration of 50 µM has been reported to induce a sleep-like state in larval zebrafish, which cannot be further enhanced with higher concentrations ^83^. Thus our findings are consistent with the activation of a sleep-like state and reduced vigilance. The violet light-induced heart rate increase in the *opn4a* (atg and spl) morphants was not further decreased when morphants were treated with 50 µM melatonin (DMSO: 3 dpf: 11.63±1.85 bpm vs. melatonin: 13.67±1.87 bpm; 4 dpf: DMSO: 15.34±1.56 bpm vs. melatonin: 13.80±2.05 bpm; s.e.m) (Fig. 4a), suggesting that the remaining response must be mediated by alternative systems that are not modulated by melatonin.

We next analysed the heart rate increase, locomotor reaction times and pectoral fin beating at night-time Zeitgeber (ZT) 19, which corresponds to a period in larval zebrafish in which melatonin levels are high and locomotor activity is most drastically reduced ^83^. We did observe a slight attenuation in the violet light-induced heart rate increase (ZT19: 25.85±2.66 bpm vs. ZT4: 28.46±1.50 bpm; s.e.m.), this attenuation however was not significant (Fig. 4b). Similarly, we did not observe any differences in the locomotor reaction times at night (ZT19) compared to day (ZT4) when assessed in violet light (ZT19: 52.73±15.79 s; s.e.m.), UV light (ZT19: 11.14±3.36 s; s.e.m.), UV/VL (ZT19: 6.54±1.93 s; s.e.m.), and blue light (ZT19: 180±0.0 s; s.e.m.)(Supplementary Fig. 7d compared to 7b). We did however observe that at night-time the onset of pectoral fin beating in violet light was not significantly faster (ZT19: 6.33±1.22 s; s.e.m.) as compared with UV light (ZT19: 9.33±3.41 s; s.e.m.)(Supplementary Fig. 7e), unlike during the daytime (Fig. 3e). Unexpectedly, also blue light elicited fin beating in some of the animals unlike during the daytime, reflected by the slight decrease in onset time (ZT19: 89.17±21.34 s) (Supplemental Fig. 7e).

In addition to these experiments, we also wanted to test whether a reduction in melatonin levels through melatonin receptor blocking and pineal gland ablation would have an effect on the heart rate induced by violet light. First, we pharmacologically inhibited melatonin receptors using the MT1/2 antagonist Luzindole ^85, 86^, which was shown to inhibit melatonin-mediated sleep induction in larval zebrafish ^83^. We expected to see a further enhancement of the heart rate increase induced by violet light. Six MT1 related genes have been identified in zebrafish but no MT2 related gene. To determine the effects of Luzindole on the violet light-induced heart rate increase, we first assessed the dose-response relationship. Treatment of wildtype animals with Luzindole concentrations ranging between 0.1 and 10 µM showed a small, but insignificant increase in the heart rate when exposed to violet light (DMSO: 29.8±2.24 bpm vs. 0.1 µM Luz: 36.67±2.66 bpm; 10 µM Luz: 37.0±3.0 bpm (Supplementary Fig. 9). Unexpectedly, a significant attenuation was observed when larvae were treated with concentrations of 100 µM (Luz: 19.6±4.06 bpm; s.e.m). We next assessed the effects of this antagonist in the *opn4a* morphants using 10 µM, which was within the range that slightly increased the heart rate. When comparing the heart rates between control and *opn4a* morphants (atg and splice), we did not observe any significant changes at 3 dpf (DMSO: 10.36±1.77 bpm vs. Luzindole: 15.16±2.21 bpm) but there was a small enhancement also in the morphants at 4 dpf (DMSO: 16.14±1.56 bpm vs. Luzindole 21.27±1.88 bpm; s.e.m) (Fig. 4a), which could be explained by the activation of alternative systems.

Next, we genetically ablated PG neurons in transgenic Tg(*tph2:NfsB-mCherry*)^y227^ larvae that express NTR under the serotonergic neuron-specific *tph2* promoter ^56^ (Fig. 4c and c’). This showed that the violet light-induced heart rate increase was not significantly different between the ablated and unablated controls at 3 and 4 dpf (3dpf: DMSO: 42.68±0.33 bpm vs. MTZ: 37.09±1.30 bpm; 4 dpf: DMSO: 47.64±2.90 bpm vs. MTZ: 39.82±4.23 bpm; s.e.m)(Fig. 4d). We observed similar effects after laser mediated and surgical ablation of fluorescent PG neurons (not shown). Taken together, these findings suggest that melatonin can modulate the violet light-induced heart rate increase but because these effects were insignificant at ZT19 when melatonin levels are highest, this hormone does not appear to have a major role in the response under physiological conditions.

### Violet light stimulates alertness-like responses through sympathetic activation

In mammals, RGCs project to a range of non-visual areas in the brain which balance light-dependent responses via sympathetic and parasympathetic output to the peripheral organs through separate preautonomic neurons ^40, 87, 88^. Light can activate the heart rate through the autonomic nervous system ^89^, and thus violet light could either increase the heart rate through sympathetic activation or parasympathetic deactivation. To assess which of the two systems mediates the heart rate increase, we first wanted to confirm that innervation of the heart is required for the increase induced by violet light. For this, we prepared heart explants of 3 dpf zebrafish larvae and compared their response to healthy siblings (Fig. 5a). Whereas the heart rate increase was significantly increased in the *in vivo* controls (30±5.62 bpm, s.d.), the increase was completely abolished in the explanted hearts (−7.8±2.22 bpm, s.d.). This finding suggests that the violet light response is controlled by autonomic innervation. To further validate this, we pharmacologically interfered with autonomic nerve function. Parasympathetic activity reduces the heart rate through the release of acetylcholine and the activation of muscarinic receptors. In contrast, sympathetic stimulation releases norepinephrine and adrenaline, which through β-adrenergic receptor (β-AR) activation induces cardiac muscle contraction and a heart rate increase. We first constitutively activated muscarinic receptors with the agonist carbachol to stimulate parasympathetic nerve fibres and a heart rate decrease (DMSO baseline: 90.45±4.92 bpm vs. carbachol baseline: 64.95±9.90 bpm; s.e.m). Nevertheless, violet light was able to overcome this reduction, leading to a heart rate increase above the DMSO control larvae (DMSO: 30.74±2.07 bpm vs. carbachol: 49.55±9.62 bpm; s.e.m)(Fig. 5b). We next antagonised sympathetic β-AR receptor function with propranolol, which also led to a decreased baseline heart rate (DMSO baseline: 102.6±3.59 bpm vs. propranolol baseline: 67.93±8.64 bpm; s.e.m). In contrast to carbachol, however, we observed a significantly attenuated violet light response compared with the untreated controls (DMSO: 32.70±2.92 bpm vs. propranolol: 18.69±4.16 bpm; s.e.m). Despite this reduction, we observed a residual heart rate increase, which could be due to the lower affinity of propranolol for β-ARs in zebrafish larvae ^90^. A previous report showed that the knockdown of *β1/β*2a-ARs in addition to propranolol treatment drastically reduces the heart rate in zebrafish larvae ^90^, which prompted us to use a similar approach. Not only was the baseline heart rate significantly decreased in the propranolol-treated double morphants (DMSO *β1/β2*a-AR baseline: 101.0±3.46 bpm vs. *β1/β2*a-AR+propranolol baseline: 33.78± 7.34 bpm; s.e.m), but the violet light response was also further attenuated (DMSO β1/β2a-AR: 22.0±4.06 bpm vs. *β1/β2*a-AR+propranolol: 8.22±2.41 bpm; s.e.m)(Supplementary Movie S6). These findings demonstrate that activation of the sympathetic nervous system is essential for the violet light-induced heart rate increase.

To further corroborate that the heart rate and locomotor responses are both dependent on the same neuronal circuits, we also analysed locomotor reaction times in propranolol-treated larvae. Although these larvae were touch responsive (Supplementary Movie S7), indicating that muscles are not paralyzed, they did not exhibit any locomotor response when exposed to violet light (Fig. 5c). Interestingly, also the UV light responses were absent, suggesting that sympathetic neuronal circuits control both *opn4a*-dependent and independent light-induced locomotor responses.

## Discussion

Light is well known to influence circadian rhythm and sleep behaviour but the mechanisms by which specific wavelengths of light influence alertness have not been extensively studied to date. Our findings in larval zebrafish suggest that violet light stimulates physiological responses that are reminiscent of alertness, leading to increased heart rate, locomotor activity, and pectoral fin beating, the latter of which has been proposed to increase oxygen supply in zebrafish larvae. Our findings in zebrafish fit well with observations in humans where a quicker and more shallow respiration has been associated with high states of alertness ^78^. Alertness in mammals has also been described to depend on sympathetic activity. For example, in alert goats, cerebral blood flow, heart rate, and arterial blood pressure have been associated with increased adrenergic and cholinergic activity ^91^. Also, in humans wakefulness is correlated with greater sympathetic activity, whereas sleep onset and the progression to deeper sleep stages is associated with a shift toward greater parasympathetic activity ^92^. Short-wavelength light has been shown to heighten alertness in humans with violet and blue light having different effects. Blue light has more significant effects on plasma melatonin reduction as compared to violet light. For example, a comparison between individuals exposed to 420 and 460 nm light at equal photon doses of 1.21 × 10^13^ photons/cm^2^ of retinal irradiance for a 90-minute period starting at midnight showed that the mean plasma melatonin reduction was 2-fold higher in blue light as compared with violet light ^51^. Blue light also more strongly increased the activity of brain centres that have been implicated in cognitive functions, such as in the left middle frontal gyrus and left thalamus, as compared with violet light at 430 nm in individuals performing a working memory task in fMRI ^42^. Yet, a study reported that violet light at 420 nm has more acute effects on subjective alertness than blue light at 470 nm ^50^. In this study, exposure to 470 nm light initially led to lower subjective alertness ratings, which only began to increase after 2 hours. In contrast, individuals exposed to 420 nm light remained alert throughout a 4-hour time period. Consistent with this are our findings, which suggest a similar acute alerting effect for violet light in larval zebrafish. Violet light induced a strong heart rate increase within only 2 seconds, whereas blue light increased the heart rate more moderately and with slower kinetics. Moreover, violet light also induced locomotor responses and pectoral fin beating, whereas blue light did not elicit these responses within the 180 sec of our analyses, suggesting that they may be delayed similar to the heart rate response. It will be interesting to further investigate in the future the longer-term effects of blue light also on these responses, as this will help to further dissect the role of short-wavelength violet and blue light in acute versus sustained alertness.

We discovered that the violet light-induced alertness-like responses require the photopigment Opn4a. Zebrafish possess five melanopsin genes, which are distinct in terms of their evolutionary origin and expression patterns. The *opn4x* genes (*opn4xa* and *opn4xb*) are more closely related to *Xenopus* melanopsin, whereas the *opn4m* genes (*opn4a* and *opn4*b) are more similar to mammals ^31, 55^. A fifth member, *opn4*.1, has been suggested to be of retrogene origin, but has been found to be functional in heterologous cell culture studies, with an absorbance peak at 403 nm in the violet spectrum ^33^. While circadian entrainment and pupillary light reflex have been proposed to use a peak melanopsin action spectra around 480 nm ^38, 93^, conflicting data regarding the peak absorbance of mammalian melanopsin exist, suggesting that Opn4 also has an action spectrum that is close to 420 nm. For example, Do and colleagues suggest that melanopsin in mice exists as a blue (473 nm) sensitive photopigment but also as a violet-shifted photopigment. It was suggested that it exists in a tri-stable state: blue, violet and silent, with green wavelengths having the ability to shut off the light response (silent state) ^53^. Also, Newman and colleagues demonstrated that the maximal action spectra of heterologous expressed mouse melanopsin in COS cells is at 424 nm when re-constituted with 11-cis retinal ^52^. It was suggested that the melanopsin spectrum could be shifted in COS cells by interactions with endogenous proteins or that melanopsin uses an unusual chromophore *in vivo*. Although their finding was unexpected, our study and the study by Matos-Cruz and colleagues showing violet light sensitivity for zebrafish Opn4.1 demonstrate that, at least in zebrafish, melanopsin can be activated by violet light. It will remain to be explored how the violet light specificity could be achieved. There are several possibilities: a zebrafish-specific shift of melanopsins to the violet spectrum, which may have been caused by mutations in spectral tuning sites leading to differences in the absorbance peaks ^94^. Zebrafish melanopsin homologues may have evolved with different wavelength specificities, with each constituting a different subclass, similar to different ipRGC subclasses in mammals where M1 cells subdivide into brn3b-positive and brn3b-negative cells and project into different areas of the brain, whereas M4 cells project predominately to the lateral geniculate nucleus (LGN) ^36, 88^. Opn4a-expressing cells could also integrate their own melanopsin response with that from other photoreceptors. Finally, signals could be integrated in different brain regions to achieve wavelength specificity. Further analyses of the precise neuronal circuits and molecular mechanisms underlying these responses in zebrafish will help to answer these questions.

We have chosen to analyse *opn4a* in our studies because of its close homology (~70 %) within the core region to human *OPN4* ^55^. However, unlike *opn4xa* that is expressed in RGCs, *opn4a* is found in the inner nuclear layer ^33^ and also in extra-ocular deep brain nuclei ^56^. The redundancy of melanopsin genes and their different expression patterns could be the reason why our ablation studies, including optical nerve transection, RGC ablation, as well as the morpholino knockdown studies, resulted in a remaining response to violet light. Alternatively, different systems that cooperatively mediate light responses may be present. For example, Dolgonos and colleagues reported that reflex blinks in response to bright light in anesthetized rats with an optic nerve transection are mediated by a light-induced increase of trigeminal activity ^95^. Although we observed that zebrafish somatosensory neurons are not required for the violet light response, the small residual violet light effect after optic nerve transection suggests that these neurons could potentially be activated by light via secondary mechanisms either through Opn4a or another light responsive protein. This redundancy may be critical to ensure that body homeostasis is maintained when one system fails. Our experiments ruled out temperature as a factor for the remaining increase since temperature changes are highest in green light whereas the heart rate is not significantly increased, even at the highest energy output level. We also ruled out that the zebrafish blue opsin Opn1sw2 with a demonstrated peak absorbance of 416 nm ^71^ contributes to the heart rate increase. Alternatively, the blue/violet sensitive cryptochromes, Cry 1 and Cry 2 could participate in the remaining response. *Cry1*/2 null mice exhibit accelerated and delayed free-running periodicity of locomotor activity, respectively ^96^. Both genes show rhythmic expression patterns in zebrafish and mice, and in zebrafish are moreover expressed in extra-ocular tissues ^23, 26^. It will therefore be interesting to determine the contributions of these alternative photopigments or other light-dependent neuronal systems to the violet light-induced responses in zebrafish.

Our data supports the notion that violet light may serve to enhance locomotor reaction time rather than triggering a response, which is consistent with alertness as defined by vigilance, readiness, or caution. The fact that UV light has stronger effects on the locomotor response but is less effective in stimulating the heart rate and pectoral fin beating, and that violet light responses depend on Opn4a, shows that the neuronal circuits leading to these responses differ between UV and violet light (Fig. 5d). Moreover, knockdown of *opn4a* delays locomotor responses in UV/VL, which are more similar to UV light responses, indicating that they are distinct responses. The precise networks and underlying molecular mechanisms for UV light-induced locomotor responses are not yet established but also depend on activation of the sympathetic nervous system. In *C. elegans*, similarly to zebrafish, locomotion is maximally accelerated by UV light, which depends on LITE-1, a novel ultraviolet light receptor that acts in neurons and is a member of the invertebrate Gustatory receptor (Gr) family ^97^. It has been suggested that the UV light-induced locomotor response may help escape lethal doses of UV light emitted from the sun ^97^. Since zebrafish are surface swimmers ^71^ they may have developed a similar mechanism to escape from lethal UV doses. Interestingly, while *lite*-1 mutants have significantly lower locomotion rates than wildtype controls, they also do not completely abolish the UV light response, suggesting that also in *C. elegans* multiple light-induced pathways are likely activated in parallel ^97^.

In mammals, blue light mediated activation of melanopsin stimulates circadian and non-circadian responses, such as photoentrainment and sleep-wake states, as well as pupil constriction, masking responses, and nociception (reviewed in ^88^). While our study suggests that in zebrafish, the melanopsin homologue Opn4a can be activated by violet light, further *in vivo* characterizations are necessary in mammals to disprove or prove a role for violet light in melanopsin activation. It is possible that alertness induced by violet light has evolved in diurnal species and therefore these studies may be more difficult to assess in nocturnal animal models. Additional studies in zebrafish will help to further validate zebrafish as a suitable model for alertness induced by short-wavelength light. Because larval zebrafish are optimal to perform rapid pharmacological screens, this model could facilitate the development of treatments for light-dependent human pathologies.

## Author contributions

J.E.C. and S.R. designed and performed the experiments and analysed the data. S.R. prepared the manuscript. S.R., J.E.C., T.S.L., C.B., and A.M.C. performed the experiments and analysed the data. A.M. assisted in data preparation and interpretation of the results.

## Acknowledgments

We thank Drs. Victoria L. Revell, Phyllis R. Robinson, Enrico Nasi, Stephan C. Neuhauss, Andrew L. Harris, Alvaro Sagasti and Georgeann S. Sack for their insight and critical reading of the manuscript. We further thank Dr. Alvaro Sagasti for providing resources in his lab. We are grateful to Drs. Marnie E. Halpern and Harold A. Burgess for kindly providing zebrafish pineal gland lines. We also thank Elizabeth A. Brochu for experimental assistance, and the MDI Biological Laboratory animal core service for zebrafish care. Research reported in this publication was supported by Institutional Development Awards (IDeA) from the National Institute of General Medical Sciences of the National Institutes of Health under grant number 1P20GM104318 and P20GM0103423, an institutional award from the US Department of Army (USAMRMC) under grant number W81XWH-BAA, and by the Albert and Ellen Grass foundation. We thank Jessie A. Rottersman for her contributions.

