## Supplemental File for "Short-wavelength violet light (420nm) stimulates melanopsin-dependent acute alertness responses in zebrafish"

**Supplementary Figures, Movies, Methods and References**

**Supplementary Figure Legends**

**Supplementary Figure 1**

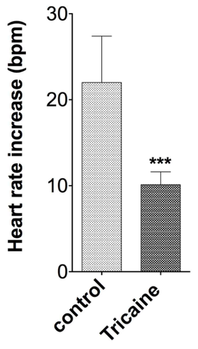


**Supplementary Figure 1. Tricaine attenuates the violet light–induced heart rate response**

Zebrafish larvae anaesthetised in Tricaine and exposed to violet light show a significantly attenuated heart rate response to violet light compared with the untreated control larvae (8 animals per group, s.e.m).

Statistical comparisons were made using the Student's t-test. ****P<0.001*

**Supplementary Figure 2**

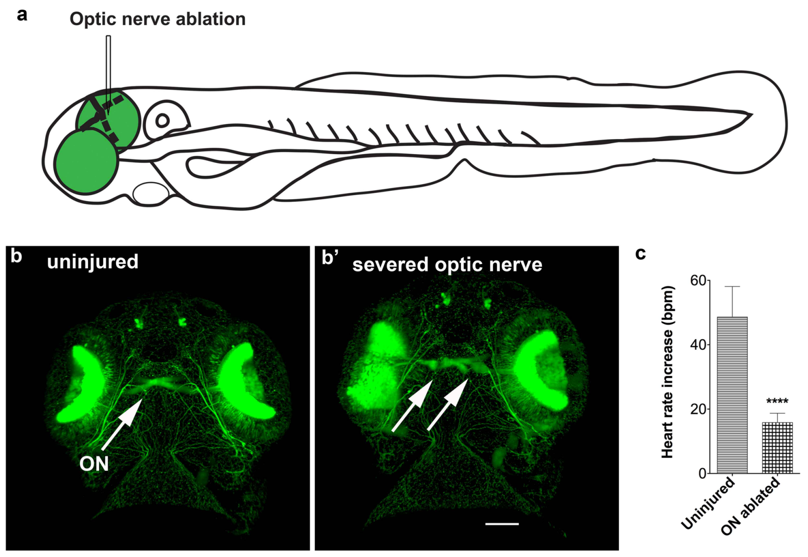


**Supplementary Figure 2. Optic nerve ablation attenuates the violet light–induced heart rate response**

(**a**) Scheme of optic nerve ablation in larval zebrafish (see Materials and Methods for a detailed description).

(**b, b'**) Optic nerves (ON, arrows) are shown before (b) and after (b') transection in a transgenic Tg(*isl2b*:GFP) larva. Scale bar: 100 µm.

(**c**) Optic nerve ablation attenuates the violet light-induced heart rate increase (6-8 animals per group, s.d.)

Statistical comparisons were made using the Student's t-test. *****P<0.0001.*

**Supplementary Figure 3**

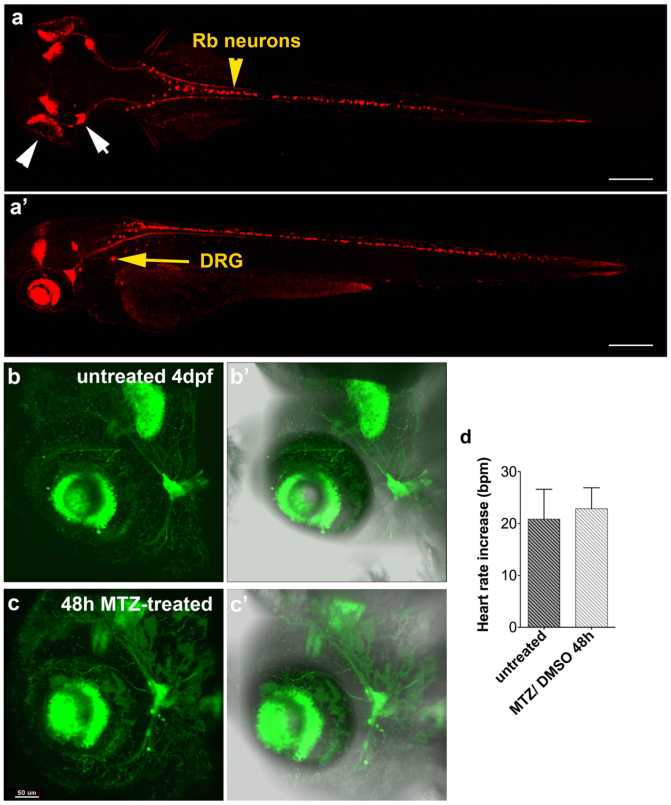


**Supplementary Figure 3. Tg(*isl2b*:*nfsB-mCherry*) larvae with *nfsB*-mCherry expression in sensory neurons and MTZ ablation in Tg(*isl2b:GFP*) larvae**

(**a, a'**) Dorsal (a) and lateral (a') view of *nfsB*-mCherry expression in 3 dpf larva, revealing fluorescence in retinal ganglion cells (RGCs; white arrowhead), trigeminal ganglia in the head (white arrow), Rohon-beard neurons (Rb neurons, yellow arrowhead), and dorsal root ganglion neurons (DRG, yellow arrow). Scale bars: 250 µm.

(**b, b', c, c'**) DMSO (b, b') and MTZ (c, c') treatments in transgenic Tg(*isl2b*:GFP) larvae. The fluorescent pattern in the MTZ-treated larva appears morphologically similar to the DMSO-treated control larva. Scale bar: 50 µm.

(**d**) The heart rate increase is also similar between the groups (7 animals per group, s.d.)

Statistical comparisons were made using the Student's t-test.

**Supplementary Figure 4**

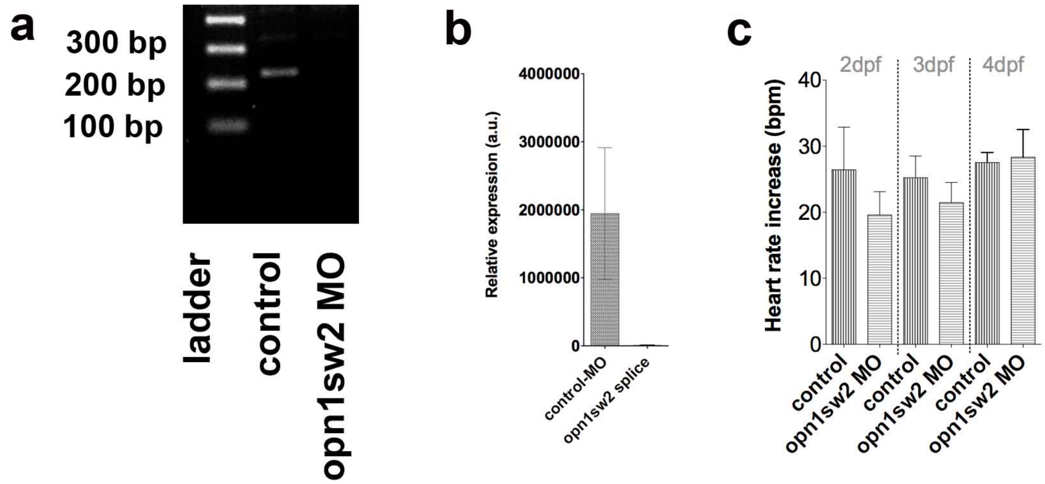


**Supplementary Figure 4. The violet light sensitive photopigment *opn1sw2* does not mediate the heart rate increase**

(**a**) RT-PCR showing nonsense-mediated RNA decay in the morpholino-injected larvae. The predicted fragment size for wildtype *opn1sw2* is 222 bp and for morphant *opn1sw2* is 307 bp.

(**b**) Quantitative PCR confirms *opn1sw2* mRNA degradation.

(**c**) Knockdown of *opn1sw2* does not significantly alter the violet light–induced heart rate increase at 2, 3 and 4 dpf. (4-46 animals per group, s.e.m.)

Statistical comparisons were made using Student’s t-test (b) and One-Way ANOAVA (c).

*P*<0.05*

### Supplementary Figure 5

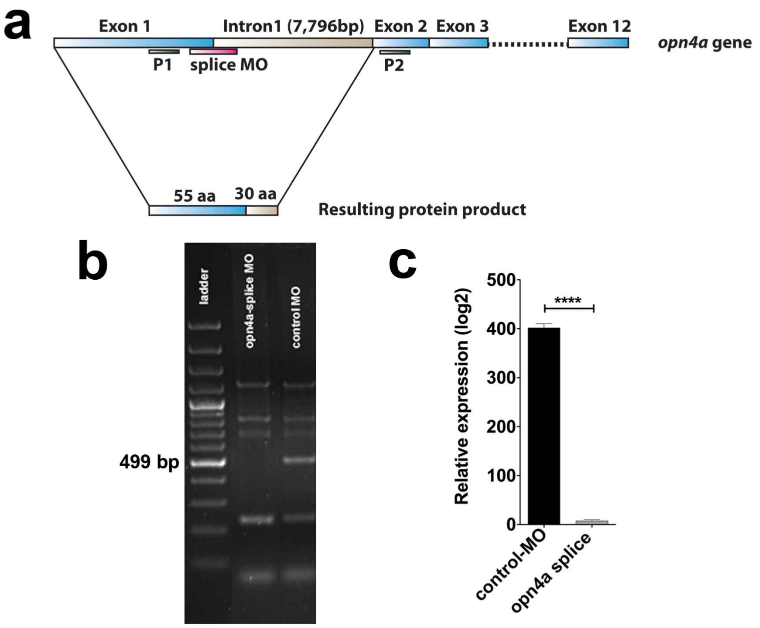


**Supplementary Figure 5. *opn4a* splice morpholino design and validation**

(**a**) The *opn4a* splice morpholino binds to the exon1/ intron 1 boundary, leading to the insertion of intron 1 and a truncation of the resulting protein. Primers were designed to anneal to exon 1 and exon 2.

(**b**) RT-PCR showing the absence of a PCR product at 499 bp in the *opn4a* morphant but not control morphant sample when the PCR extension time was adjusted to the wildtype product.

(**c**) Quantitative PCR confirming the absence of a product.

Statistical comparisons were made to using the Student's t-test. *******$P<0.0001$***

### Supplementary Figure 6

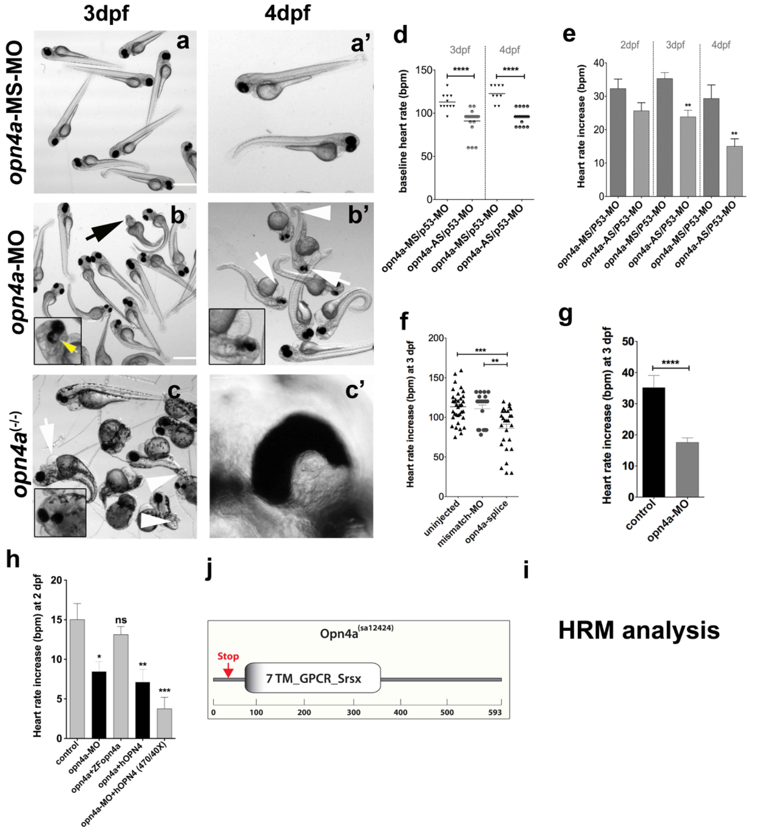


**Supplementary Figure 6. *opn4a* knockdown and violet light-induced heart rate increase in *opn4a* splice morphants with and without co-injection of zebrafish *opn4a* and human *opn4* mRNA**

(**a-c, a'-c'**) Unlike *opn4a*-missense MO-injected control larvae (a, a'), *opn4a*-antisense morphants (b, b') and *opn4a*$^{(sa12424)}$ mutants (c, c') develop truncated tails (white arrowheads), smaller eyes (black arrow, insets in b, c), heart edema (white arrows) and depigmented ventral retinae (insets in b, c). The extent of these mutant phenotypes is variable. Scale bars: 750 µm. (**d**) Comparison of the baseline heart rate between mismatch and antisense *opn4a*-ATG morphants co-injected with a *p53*-MO at 3 and 4 dpf. The baseline heart rate in the morphants is significantly lower than in the two controls (19-35 animals per group, s.e.m).

(**e**) Heart rate analysis at 2, 3 and 4 dpf shows a significant attenuation of the violet light-induced-induced heart rate increase for the *opn4a* splice morphants co-injected with a *p53*-MO (24-54 animals per group, s.e.m).

(**f**) Heart rate analysis at 2 dpf reveals a significant attenuation of the violet light-induced-induced heart rate increase for *opn4a*-splice morphants, which can be rescued by co-injection of zebrafish *opn4a* mRNA but not when human *OPN4* mRNA is co-injected. Human OPN4 mRNA also does not mediate a response when stimulated with blue light (470/40X filter) (6-16 animals per group, s.e.m.).

Asterisks above columns represent comparisons to the first column in each group. Brackets indicate alternative comparisons. Statistical comparisons were made using Student's t-test (e) and One-Way ANOAVA (d, f). **P<0.05, **P<0.01, ***P<0.001, ****P<0.0001; Abbreviations: mismatch, MS; antisense, AS; hOPN4, human OPN4.*

### Supplementary Figure 7

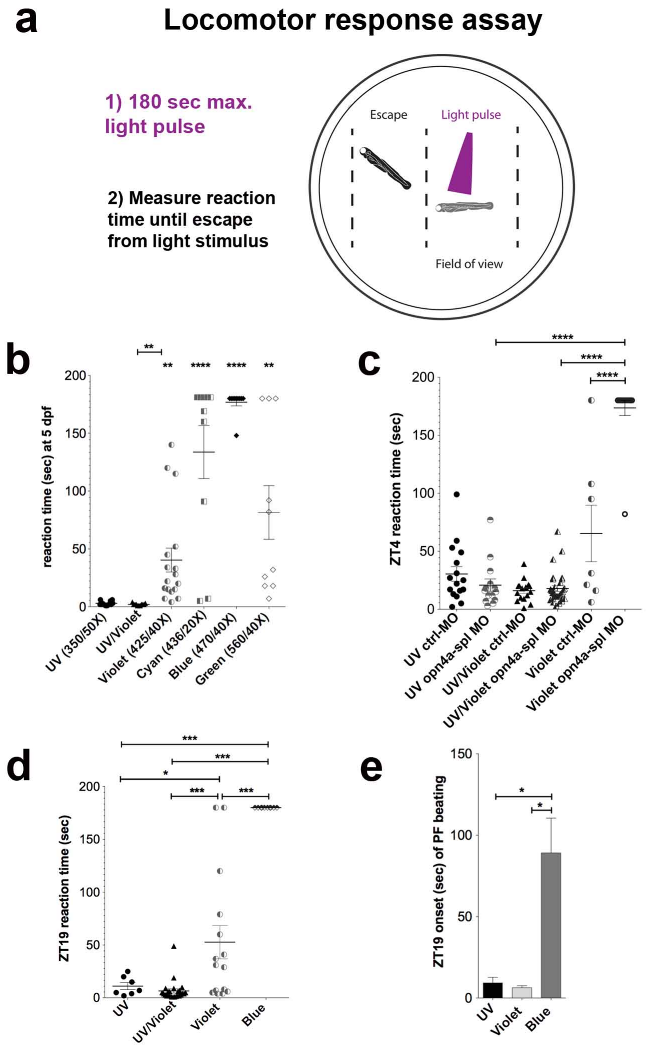


**Supplementary Figure 7. Locomotor response and pectoral fin beating analyses at 5 dpf, in *opn4a* splice morphants, and at ZT19**

(**a**) Scheme of locomotor response assay. The time until the larvae escaped from the light stimulus was measured for up to 180 seconds.

(**b**) The locomotor response at 5 dpf for UV, UV/violet, cyan, and green light is shown. Cyan and blue light rarely elicit a locomotor response within the measured time (n=10-17 animals per group, s.e.m).

(**c**) Locomotor response in the *opn4a*-splice morphants at 3 dpf. Violet light does not elicit a locomotor response in the morphants unlike UV light and UV/VL (n=7-32 animals per group, s.e.m).

(**d**) Zeitgeber (ZT) 19 locomotor reaction time assessed in wildtype larvae in UV, UV/VL, violet and blue light. The response is comparable to daylight reaction times (7-25 animals per group, s.e.m.).

(**e**) Pectoral fin beating onset in UV, violet and blue light measured at ZT19 (6-13 animals per group, s.e.m.).

Asterisks above columns represent comparisons to the first column. Brackets indicate alternative comparisons. Statistical comparisons were made using One-Way ANOAVA. *P*<0.05, **P<0.01, ***P<0.001, ****P<0.0001*

**Supplementary Figure 8**

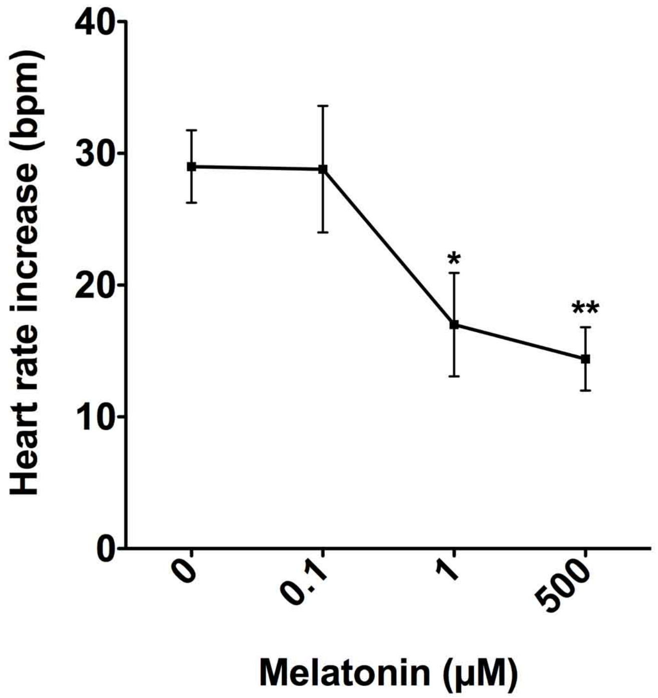


**Supplementary Figure 8. Dose-response curves for melatonin**

Pharmacological melatonin treatment significantly attenuates the violet light-induced heart rate increase at concentrations at and above 1 μM (5 animals per group, s.e.m.).

Asterisks above treatment groups represent comparisons to the untreated group. Statistical comparisons were made using One-Way ANOAVA. *P*<0.05, **P<0.01*

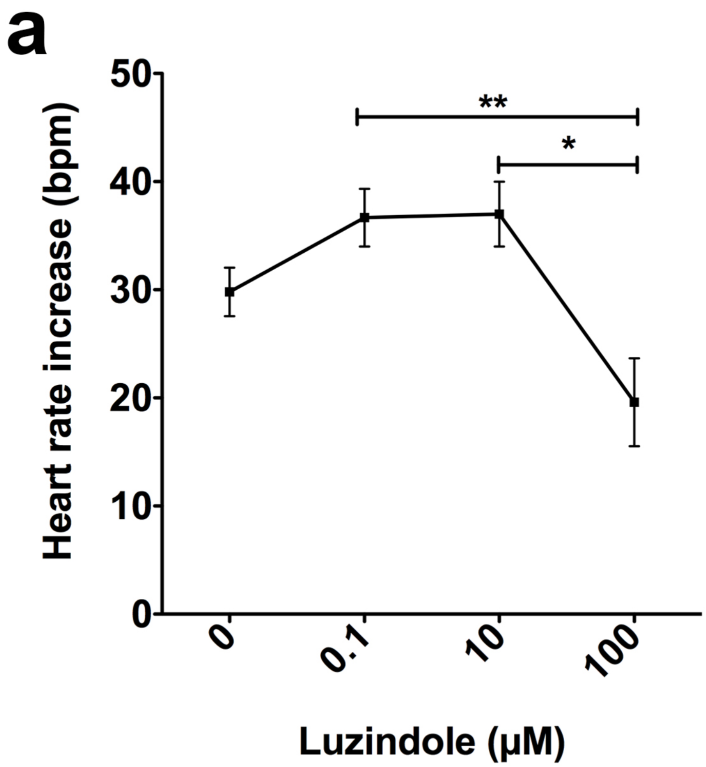


**Supplementary Figure 9. Dose-response curves for Luzindole**

Treatment of wildtype animals with various Luzindole concentrations shows a small increase in the violet light-induced heart rate at 0.1µM and a significant attenuation in the heart rate increase induced by violet light at 100 µM. Statistical comparisons were made using One-Way ANOAVA. *P*<0.05, **P<0.01*

**Supplemental Movie Legends**

**Supplementary Movie 1: Violet light-induced heart rate increase and recovery.** Shown is a larva exposed for 5 sec to 1) bright light, 2) bright light + violet light, and 3) bright light. Transitions are indicated in the movie. The heart rate rapidly increases in violet light and decreases back to pre-exposure levels upon termination of the violet light exposure.

**Supplementary Movie 2: Heartbeat in an *opn4a* morphant larva prior to and during violet light exposure.** Shown is a larva exposed for 5 sec to 1) bright light and 2) bright light + violet light. The heart rate remains at a similar rate throughout the violet light exposure.

**Supplementary Movie 3: Pectoral fin beating increases during violet light exposure.** Shown is a larva with increased pectoral fin beating during violet light exposure.

**Supplementary Movie 4: Pectoral fin beating response in UV light.** Shown is a larva exposed to bright light for 10 sec and subsequently to UV light for 2 min, which shows a delayed pectoral fin beating onset and a lower rate of beating.

**Supplementary Movie 5: Blue light does not trigger pectoral fin beating.** Shown is a larva exposed to bright light for 10 sec and blue light for 2 min. Blue light does not elicit pectoral fin beating during the time of the exposure.

**Supplementary Movie 6: Heartbeat in a *$\beta$1/2a-AR* double-morphant larva treated with propranolol.** Shown is a propranolol-treated morphant larva exposed to 5 sec of bright light and subsequently to 5 sec of bright light plus violet light. The heart rate remains similar prior to and during violet light exposure.

**Supplementary Movie 7: Locomotor response of a propranolol-treated larva.** Shown is a larva propranolol-treated larva responding to a touch stimulus, which suggests that the muscles are not paralysed by propranolol.

### Supplementary Methods

#### Zebrafish husbandry

Raising, spawning, and maintenance of zebrafish was performed according to NIH guidelines. Embryos up to the larval stages were raised at 28°C and maintained in 0.03 % Instant Ocean salt solution (fish water). The following wildtype strains were analysed: AB, Tuebingen (Tu), Tu x Tupfel long fin (Tu/TL), and Nacre. Transgenic Tg(*isl2b*:GFP) *(11)* fish were obtained from Dr. Alvaro Sagasti's lab (courtesy of Chi-Bin Chien, formerly at the University of Utah). Embryos and larvae were kept on a 14:10 h light/dark (LD) cycle.

#### Fluorescent filters

The following fluorescence filters were used for the wavelength-specific analyses:

<u>Kramer Scientific, LLC (Amesbury, MA, USA):</u>

350/50X; 390/70X; 470/40X; 535/50X; D560/40X

Chroma Technology Corporation (Bellows Falls, VT, USA):

405/10X; 425/40X; 436/20X; D545/25X

**Pharmacological administrations**

Larvae were treated for 2 h prior to the heart rate measurements with carbachol, propranolol, melatonin and Luzindole as indicated in the text (all reagents were obtained from Sigma-Aldrich). For the ablation experiments, transgenic Tg(*isl2:nfsB-mCherry*) and Tg(*isl2:GFP*)[1] larvae were treated at 48 hpf for either 24 or 48 h with 13 mM MTZ/0.5 % DMSO. The pineal glands were ablated by treating transgenic Tg(*tph2*:NfsB-mCherry)$^{y227}$ larvae [2] at 2 and 3 dpf for 24 h with 10 mM MTZ/0.5 % DMSO.

**Cloning of zebrafish *opn4a***

Primers targeting the 5' and 3' UTR of zebrafish *opn4a* with the following sequences were designed: Fwd: 5'- AACGCTTCTCTCACCG -3' and Rev: 5'- CCTCCTCTCCCTCTGTTT -3'. *opn4a* was amplified from cDNA that was generated from 4 dpf Nacre fish using the proof-reading enzyme primeSTAR GXL (Clontech). The amplified cDNA was cloned into the pCR-Blunt II-TOPO vector (Life Technologies) and messenger RNA was generated using the Ambion T7 Message Machine kit (Life Technologies). The mRNA (100 ng/µl) was co-injected with 3-10 ng of the *opn4a* splice morpholino.

**Preparation of human *OPN4* mRNA**

Human *OPN4* cDNA was obtained from Open Biosystems (Fisher Scientific, Pittsburgh, PA, USA), and mRNA was prepared using the Sp6 Message Machine kit (Ambion – Life Technologies, Green Island, NY, USA). One hundred pg of mRNA was injected into 1-cell stage embryos together with 3-10 ng of *opn4a* morpholino.

### Morpholino sequences

Morpholino sequences are shown in Tables 1 and 2.

Table 1: Designed morpholinos.

| **Morpholino name** | **5' – 3' sequence** |
|---|---|
| *opn4a*-ATG mismatch | GCGCCTCCCCTGATCATCATGACCA |
| *opn4a*-ATG antisense | GCGGCTCCGCTCATCATCATCACGA |
| *Opn4a*-splice antisense | ATCTAGGACTCACAGATGTAGTAGT |
| *opn1 sw2*-ATG antisense | GTTTGCTGTTGTTGCTTCATCTTGC |
| *opn1 sw2*-ATG mismatch | GTATCCTGTTCTTCCTTCATCTTCC |
| *Opn1sw2*-splice antisense | GGACGCATGTTACAATGTACCTCCA |

Table 2: Previously published and validated morpholinos.

| **Morpholino name** | **5' – 3' sequence** | **References** |
|---|---|---|
| *p53* antisense | GCGCCATTGCTTTGCAAGAATTG | Gene Tools, LLC, USA |
| *β1-AR* antisense | ACGGTAGCCCGTCTCCCATGATTTG | *(30)* |
| *β2a-AR* antisense | GTATTGAGGACCTTATGTTTCCCAT | *(30)* |
| *ngn1* antisense | CCATATCGGAGTATACGATCTCCAT | *(12)* |
| *Standard Control* | CCT CTT ACC TCA GTT ACA ATT TAT A | Gene Tools, LLC, USA |

Morpholinos were dissolved in 100 μl or 300 μl of double-distilled water to make 3 and 1 mM stock solutions, respectively, and stored at 4°C until use.

**Reverse Transcription and quantitative PCR for *opn1sw2* and opn4a splice morpholino validation**

Fifteen larvae at 2 dpf were pooled prior to RNA isolation using the Trizol method. The mRNA was reverse transcribed using random hexamer and oligo dT primers. RT-PCR was performed using the following primers for *opn1sw2* annealing in exon1 (fwd: AAT CGA GGG CTT CAC TTC AA) and exon 3 (rev: CAC TGC AAG CCC TCA GGT AT). Amplification of *opn1sw2* was predicted to yield a 307 bp fragment due to the inclusion of intron1 in the morpholino-injected larvae, whereas the wildtype fragment is predicted to be 222 bp.

The following primers were designed for *opn4a,* annealing in exon 1 (fwd: CTCTCACCGGCTCAAAACTC) and exon 3 (rev: CTTTCTCTCCAAAGATCCAT). The resulting splice morpholino product is 7.8 kb larger than the wildtype product (499 bp) due to the inclusion of intron 1.

**Generation of somatosensory neuron-deficient larvae**

Embryos of the transgenic Tg(*isl2b*:GFP) strain were injected with 5 ng of *ngn1*-MO at the 1- to 4-cell stage and raised until 2 dpf. The morphants were screened under a fluorescence stereomicroscope (Zeiss Discovery II) for GFP expression in the eye and for the absence of somatosensory neurons (trigeminal ganglia, dorsal root ganglia, and Rohon-beard neurons); the morphants were analysed at 3 dpf for the effects of violet light on the heart rate.

**Optic nerve dissections**

Optic nerves were dissected with a pulled glass capillary. The capillary was carefully broken and inserted from the dorsal to the ventral side behind the eyes of the transgenic Tg(*isl2b*:GFP) larvae, which allowed for the visualisation of RGCs under a fluorescent stereomicroscope (Zeiss Discovery II). The inserted needle was moved in horizontal direction until the optic nerve was dissected. The manipulated larvae were analysed on an Olympus FluoView FV1000 confocal microscope to verify the successful nerve transection.

#### Generation of transgenic Tg(*isl2b*:*nfsB-mCherry*) fish

For genetic ablation of RGCs, we generated a transgenic Tg(*isl2b*:*nfsB-mCherry*) line harbouring the gene encoding bacterial Nitroreductase (*nfsB*) fused to mCherry, which is expressed under the *islet2b* promoter [1] in somatosensory neurons and RGCs. The expression vector was generated using the Gateway System (Life Technologies, USA). The destination vector was assembled from a 5' pDONR vector that contained the *islet2b* promoter, a middle pDONR vector that contained the *nfsB-mCherry* cassette (courtesy of Michael Parsons), and a 3' pDONR vector that contained a poly-Adenylation signal sequence. The destination vector contained Tol2 sites at the 5' and 3' ends (courtesy of Chi-Bin Chien), flanking the insert for genomic integration. Transgenic fish were generated by co-injecting 300 pg of Tol2 transposase mRNA and 50 pg of plasmid DNA into early 1-cell stage embryos. Transgenic fish were bred to homozygosity prior to neuronal ablations.

#### Heart rate analysis

For the heart rate analyses, unanesthetized larvae were mounted in 1.2 % low-melt agarose (Fluka). The larvae were then covered with fish water that was adjusted to 21°C room temperature. The following equipment was utilised for the analyses: 1) Nikon Eclipse Ti-U with a 10x eyepiece lens, 4x objective, a Prior Scientific Lumen 200 Fluorescence Illumination System, and Stanford Photonics Inc.'s Piper Control Acquisition software and an X-Cite® 120Q metal-halide lamp (120 Watt), 2) a Zeiss Discovery II fluorescence stereomicroscope with a 3.5x objective, an X-Cite® 120Q metal-halide lamp (120 Watt) equipped with Axiovision time-lapse module, and 3) a Zeiss Axioplan with a 4x EC-Plan Neofluar objective, operated with an X-Cite® 120Q metal-halide lamp (120 Watt) equipped with Axiovision time-lapse module.

The heartbeats were recorded at maximal speed (14-48 frames/sec) using the following standardised procedure: Mounted larvae were placed in bright light and subsequently additionally exposed to fluorescent light. Movies were recorded for 5 sec in bright light, followed by 5 sec in bright plus fluorescent light.  For recovery studies, an additional 5 sec post-exposure were recorded. Heartbeats were subsequently determined in each movie and the heart rate calculated, expressed as beats per minute (bpm).  The bright light intensities were dependent on the microscope settings utilized: 425/40X: 64-800 µW/cm$^2$ and for the remaining filters: 4-6 mW/cm$^{22}$. In general, the bright light intensity did not influence significantly the heart rate increase.

**Behavioural analyses**

Three or 5 dpf zebrafish larvae were assessed for locomotor activity and pectoral fin beating using a Zeiss Discovery Fluorescence Stereomicroscope equipped with a 3.5x objective and Axiovision software; a Nikon Eclipse Ti-U microscope equipped with a 10x eyepiece lens, 4x objective, and Stanford Photonics Inc.'s Piper Control Acquisition software; or a Zeiss Axioplan equipped with a 4x EC-Plan Neofluar objective and Axiovision software. Prior to analysis, locomotor activity was assessed through sensory stimulation by tapping the otoliths with a 200 µl extended-length gel pipette tip to trigger an escape response. The locomotor responses in fluorescent light were measured by placing a single larva into a Petri dish that contained 21°C fish water. Prior to the light stimulus, the larvae were kept in bright light for 30 sec, which we determined did not elicit a locomotor response. Following the initial bright light exposure, the larvae were additionally exposed to fluorescent light (UV, UV/violet, violet, cyan, blue, or green) until they escaped from the field of view or for a maximum period of 180 sec. For the pectoral fin beating analyses, the larvae were maintained for 60 sec in bright light and then additionally exposed to fluorescent light for 180 sec. The fin beats per minute and the time until onset of fin beating occurred in fluorescent light were determined. Only larvae that did not show locomotor behaviour prior to fluorescent light exposure were included in the analysis. The bright light intensities were dependent on the microscope settings utilized: 425/40X: 64 $\mu W/cm^2$ and for the remaining filters: 0-100 $\mu W/cm^{2}$[2]. Sudden bright light exposure was tested in the ability to trigger a locomotor response or pectoral fin beating but was found not to influence these responses.

**Heart explants**

Larvae at 2 dpf were anaesthetised in 0.4 mM MS-222, and one heart at a time was separated from the body by squeezing the cardiac sac on both ventrolateral sides with forceps, while simultaneously tearing off the heart from above the yolk sac in the posterior to anterior direction. The hearts were maintained in isotonic Tyrode's solution for <15 minutes prior to violet light analyses.

**Statistical analyses**

Statistical comparisons were made using Prism 5 software (GraphPad Software, Inc.). As indicated in the figures, the unpaired Student's t-test with a 95 % confidence interval was used to compare the means of two unmatched groups, assuming that the values followed a Gaussian distribution. For multiple comparison tests of three or more groups, one-way ANOVA at an alpha=0.05 (95 % confidence interval) and Tukey's multiple comparison post-tests were utilised to compare the means of each column. Significance is denoted with asterisks: *$P<0.05$, **$P<0.01$, ***$P<0.001$, ****$P<0.0001$.
